# Malayan kraits (*Bungarus candidus*) show affinity to anthropogenic structures in a human dominated landscape

**DOI:** 10.1101/2021.09.08.459477

**Authors:** Cameron Wesley Hodges, Benjamin Michael Marshall, Jacques George Hill, Colin Thomas Strine

## Abstract

Animal movement can impact human-wildlife conflict through a variety of features: increased movement can lead to greater chance of encounter, remaining still can lead to greater or lesser detection, and activity can modulate their impact on humans. Here we assess the movement and space use of the highly venomous and medically important Malayan krait (*Bungarus candidus*) on a suburban university campus. We radio-tracked 14 kraits for an average of 114 days (min: 19, max: 218), with an average of 106 fixes (min: 21, max: 229). We assessed movement pathways and activity with dynamic Brownian Bridge Movement models, evaluated patterns of revisitation to identify site fidelity, and determined whether individuals selected for particular land-use types with Integrated Step Selection Functions. Most individuals displayed some level of attraction to buildings (n = 10) and natural areas (n = 12); we identified a similar unambiguous pattern of attraction to buildings and natural areas at the population level (of our sample). Snakes remained in shelter sites for long periods of time (max = 94 days) and revisited sites on average every 15.45 days. Over 50% of fixes were within human settlements and 37.1% were associated with buildings. We found generally seasonal patterns of activity, with higher activity in wet seasons (when classes are typically in session on campus), and lower activity in the hot season (when there are a number of short breaks causing students to leave campus). These results show frequent proximity between Malayan kraits and humans at Suranaree University of Technology, Thailand; thereby, suggesting a near constant potential for human-wildlife conflict. Despite the fact that no snakebites from this species occurred at the university during our study period, substantial education and awareness training should be considered to raise awareness to ensure continued coexistence on campus.

## Introduction

Animal movements are driven by the behavior and the search to fulfil animal’s needs (Avgar et al., 2013), needs that can be met by resources: food, water, shelter, mates, and suitable nesting sites. Such movement is also modulated by simultaneous efforts to minimize predation risk (Eifler and Eifler, 2014; Liu et al., 2021), avoid inhospitable areas (Beyer et al., 2016), and conserve energy or water (Christian, Webb and Schultz, 2003). Barriers to animal movements are increasingly anthropogenic, such as impassable structures (e.g., roads, walls, and dams), as well as large expanses of developed areas or agricultural land, which is unsuitable for many species of wildlife (Brooks, 2003; Brodie et al., 2016; Berger-Tal and Saltz, 2019). To mitigate the negative consequences of anthropogenic landscape modifications, we require a deeper and broader understanding of how animals move in/through such areas. Movement strategy influences survival and ultimately fitness, in turn influencing conservation or conflict management decisions (Andreassen and Ims, 1998; Doherty and Driscoll, 2018).

Regardless of the potential negative consequences for wildlife, there are numerous examples of animals exploiting even heavily modified areas (Duarte et al., 2011; Garner et al., 2020; Bista et al., 2021).

However, such cohabitation can lead to conflicts when animal or human impacts the other’s health or access to resources. Animals living among humans can impact human health and quality of life, whether by spreading disease like rodents (Morand, Jittapalapong and Kosoy, 2015) or mosquitos (Dahmana and Mediannikov, 2020), damaging property or livelihoods (Wilson et al., 2015; Gross et al., 2021), or by directly causing injury or death (snakebite). Thus a better understanding of movement and behavior of conflict prevalent species in relation to humans can aid in developing preventative measures (Messmer, 2000). Snakebites afflict more than 2.5 million people globally, contributing annually to 81,410-137,880 deaths and >400,000 permanent injuries (including amputation, restricted mobility, blindness, and extensive scarring) to people worldwide (Chippaux, 1998; Kasturiratne et al., 2008; Gutiérrez et al., 2017; World Health Organization, 2019; Suraweera et al., 2020). In 2017 the World Health Organization declared snakebite envenomations as a highest priority neglected tropical disease (Chippaux, 2017).

Despite the clear need, and growing body of literature (Glaudas, 2021, Banes et al., 2018; Knierim et al., 2019), there is still a demand for more information on snake spatial ecology in human-dominated landscapes if we are to determine potential conflict hotspots –particularly in the tropics where snakebite is a major issue (Malhotra et al., 2021).

Malayan kraits (*Bungarus candidus*) cause life-threatening envenomings to humans across their distribution in Southeast Asia. In Thailand (one of the countries with published data on snakebite incidence in Southeast Asia), the Malayan krait has the highest mortality rate and was tied with *Calloselasma rhodostoma* for the greatest number of mortalities in one study (Looareesuwan, Viravan and Warrell, 1988; World Health Organization, 2016). Bites by kraits (*Bungarus spp.*) can be painless and often occur during the night, with some victims bitten while sleeping (Prasarnpun et al., 2005; Warrell, 2010; Tongpoo et al., 2018). It is critical to develop better strategies to prevent bites from kraits, and one of the first steps towards this is to better understand the species behavior and ecology (Malhotra et al., 2021). Presently, the literature remains limited to four studies published based on the movements, space use, or habitat use of Malayan kraits, all with a sample size of one individual each. These preliminary studies informed our theoretical framework, helping us to generate hypotheses. Three took place in the Sakaerat Biosphere Reserve, northeast Thailand: one within the dry Dipterocarp forest (Mohammadi et al., 2014), and two within neighboring agricultural areas in the transitional/buffer zone of the biosphere reserve (Crane et al., 2016; Knierim et al., 2018). The most recent publication was a focal animal study – on an individual from this study– highlighting potential conflict on a university campus (Hodges et al., 2021).

This study is the first to examine the spatial ecology of multiple Malayan kraits *Bungarus candidus* (LINNAEUS, 1758), with methods that treat positional data as inherently autocorrelated movement data (such as autocorrelated kernel density estimates). Here we use radio-telemetry to assess the space use, temporal movement patterns, and habitat selection of a population living among a complex mosaic landscape of human-modified lands (a university campus and surrounding area) in northeast Thailand. We hypothesized that telemetered *B. candidus* individuals would spend most of their time in heavily vegetated areas (e.g., forest fragments). We also hypothesized that *B. candidus* will exhibit strong site fidelity, returning to some shelter sites multiple times. We used a suite of movement driven approaches to assess these hypotheses in an empirical manner.

## Methods

### Study site

The study area covers the campus of Suranaree University of Technology (SUT) and its surrounding landscape in Muang, Nakhon Ratchasima, Thailand (14.879°N, 102.018°E; Figure 1). The university campus covers about 11.2 km^2^, and comprises a matrix of human modified lands interspersed with mixed deciduous forest fragments (at the onset of this study we identified there were 37 mixed deciduous forest fragments on campus, mean = 7.36 ±1.48 ha, range = 0.45-45.6 ha [note “±” is used for standard error throughout the text]). More than 15,000 students are enrolled at SUT (SUT Division of Planning, 2017), and there are numerous multi-story classrooms, laboratory and workshop buildings, residential housing, parking areas, eating and sports facilities, an elementary school, and a large hospital on the university campus. During the first term of the 2019 school year, 7,622 students, as well as numerous SUT staff, lived in on-campus residential areas. The landscape surrounding the university is primarily dominated by agriculture, though there are also patches of less-disturbed areas as well as several densely populated villages and suburban housing divisions among the monoculture plots of upland crops (e.g., cassava, maize, and eucalyptus).

**Figure 1.**
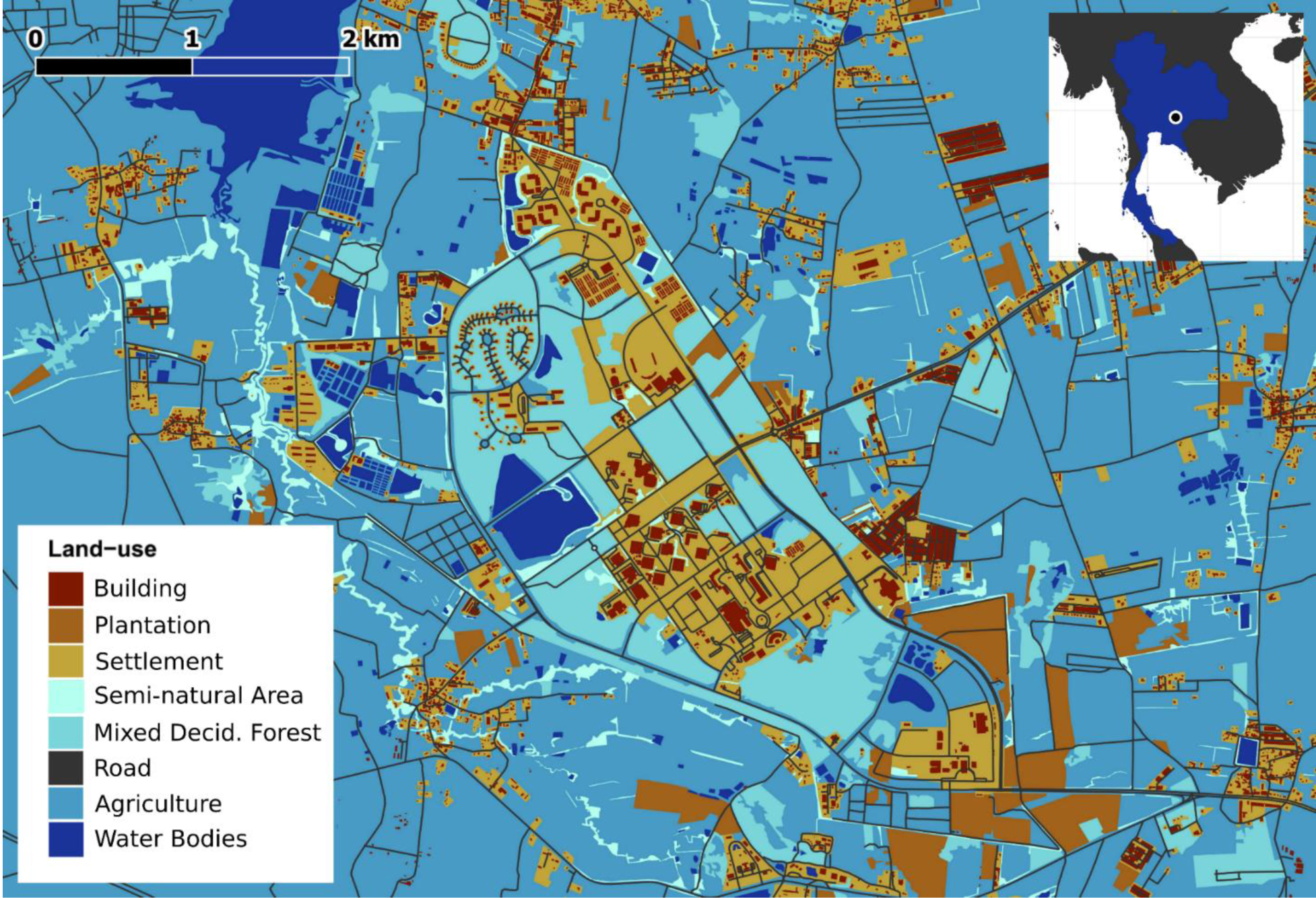
Study site map illustrating the land-use types spanning the area where the Malayan kraits (*Bungarus candidus*) were tracked in Muang Nakhon Ratchasima, Nakhon Ratchasima province, Thailand.

The study site is located within the Korat Plateau region with an altitude range of 205-285 m above sea level. Northeast Thailand has a tropical climate, and the average daily temperature from 1-1-2018 to 31- 12-2020 in Muang Nakhon Ratchasima was 28.29°C, with daily averages ranging from 19.3 to 34.1°C (average daily temperature data provided by National Oceanic and Atmospheric Administration, 2021). The region receives an average annual rainfall ranging from 1,270 to 2,000 mm (Babel et al., 2011).

There are three distinct seasons in Northeast Thailand: cold, wet, and hot, each are classified by annual changes in temperature and rainfall. Cold season is typically between mid-October and mid-February, hot season is generally from mid-February to May, while the highly unpredictable rainfall of the wet season is predominantly concentrated between the months May to October (Babel et al., 2011; Thai Meteorological Department, 2014).

Due to the representation of agriculture, semi-urban, and suburban areas with patches of more natural areas all within a relatively small area, we determined the university campus provided an ideal setting to examine how land-use features and human activity influence the movements of *B. candidus*. Additionally, past studies have indicated northeast Thailand hosts the most bites by *B. candidus* in Thailand (Looareesuwan, Viravan and Warrell, 1988; Tongpoo et al., 2018), making sites like ours ideal.

### Study animals

We opportunistically sampled Malayan kraits captured as a result of notifications from locals and ad-hoc encounters during transit due to low detectability in visual encounter surveys, in addition to those discovered through unstandardized visual encounter surveys. Upon capture, we collected morphometric data, including snout-vent length (SVL), tail length (TL), mass, and sex (Table 1, Supp. Table 1). We measured body lengths with a tape measure, measured body mass with a digital scale, and determined sex via cloacal probing, all while the snakes were anesthetized via inhaling vaporized isoflurane. We then housed individuals with an SVL >645 mm and mass >50 g in plastic boxes (with refugia and water) prior to surgical transmitter implantation by a Veterinarian from the Nakhon Ratchasima Zoo. We attempted to minimize the time snakes were in captivity awaiting implantation; however, delays arose due to the Veterinarian’s availability, the snake being mid-ecdysis, or the snake having a bolus that needed to pass through the digestive tract before implantation (n = 21 implantations, mean = 5.02 ±0.61 days, range = 0.60-13.02 days). The Nakhon Ratchasima Zoo vet implanted radio transmitters (1.8 g BD-2 or 3.6 g SB- 2 Holohil Inc, Carp, Canada) into the coelomic cavity using procedures described by Reinert and Cundall (1982), while the snake was anesthetized. We assigned each individual an ID according to sex and individual detection number (e.g., M02 = a male was the second *B. candidus* individual documented during the study). We released the implanted individuals as close as possible to their capture locations (mean = 65.31 m ±13.7 m, range = 0-226.42 m), though on six occasions we moved individuals ≥100 m because the individual came from either residential areas or a busy road (all but one were moved <155 m; see Supp. Table 2 for full details on captures, surgeries, and releases). Cameron Hodges released all snakes within 12 hours post-surgery (after nightfall), though on one occasion retained two individuals (M32 and M33) for an additional night post-surgery to avoid heavy rainfall the night of the implantation surgery. We began collecting location data the day after their release. We included all tracking data in analyses, as the animals appeared to move and behave as usual immediately following their release.

**Table 1.** Morphometric and tracking data for each telemetered *B. candidus*. Mass, snout-to-vent length (SVL), tracking start and end dates (shown as year-month-day), lag-time between location checks (hours), number of fixes, duration tracked (days), and number of relocations (i.e., moves) shown with overall averages (for all include, n = 14).

| ID | Mass<br>(g) | SVL<br>(mm) | Start | End | Lag-time (h) | Fixes | Days Tracked<br>(days) | Moves |
| --- | --- | --- | --- | --- | --- | --- | --- | --- |
| M01 | 339.5 | 1130 | 18-6-9 | 18-9-15 | 20.58 ± 1.31 | 115 | 97.77 | 34 |
| M02 | 287.7 | 1081 | 18-9-21 | 19-4-27 | 22.98 ± 0.98 | 229 | 218.27 | 58 |
| M07 | 248.4 | 1013 | 18-10-25 | 19-3-13 | 26.06 ± 2.45 | 129 | 138.98 | 23 |
| M12 | 544.3 | 1303 | 18-11-22 | 19-4-27 | 26.43 ± 2.72 | 143 | 156.35 | 23 |
| M14 | 218 | 914 | 18-12-15 | 19-2-28 | 23.99 ± 0.18 | 76 | 74.96 | 17 |
| F16 | 216.7 | 912 | 19-1-15 | 19-5-4 | 23.55 ± 0.26 | 111 | 107.92 | 14 |
| M22 | 63.6 | 650 | 19-5-1 | 19-6-19 | 23.94 ± 0.37 | 50 | 48.88 | 26 |
| M27 | 91.5 | 727 | 19-6-21 | 19-7-27 | 28.08 ± 4.65 | 31 | 35.1 | 8 |
| M28 | 91.2 | 772 | 19-7-9 | 19-10-18 | 21.06 ± 0.72 | 116 | 100.92 | 31 |
| M29 | 56.8 | 645 | 19-7-30 | 19-8-19 | 22.9 ± 1.23 | 21 | 19.08 | 3 |
| M32 | 485 | 1196 | 19-9-24 | 20-1-3 | 24.91 ± 1.02 | 98 | 100.66 | 23 |
| M33 | 176.7 | 904 | 19-9-25 | 20-3-23 | 24.39 ± 0.46 | 178 | 179.89 | 31 |
| M35 | 450 | 1113 | 19-10-17 | 20-3-4 | 24.03 ± 0.21 | 140 | 139.18 | 18 |
| M36 | 500 | 1456 | 19-12-6 | 20-2-4 | 27.64 ± 3.69 | 53 | 59.89 | 16 |
| Avg | 269.2 | 986.86 | na | na | 24.20 ± 0.41 | 106.43<br>± 15.43 | 105.56 ± 15.16 | 23.21<br>± 3.55 |

We radio-tracked 14 individuals (13 males, 1 female) between 8 June 2018 and 24 March 2020 within the SUT study site (Table 1) and classed individuals as adults if the SVL was >800 mm; thus, nine of the males were adults and four were juveniles (though two of the males had an SVL >720 mm, and therefore likely sub-adults). The single telemetered female was an adult.

Individual tracking durations varied (mean = 106.46 ±15.36 days, range = 28.5-222.77 days; Supp. Figure 1), as many individuals were lost due to unexpected premature transmitter failures (n = 5) or unsuccessful recapture efforts due to individuals sheltering under large buildings as the transmitter reached the end of its battery life (n = 4). We only recorded one confirmed mortality in the study, M01, who was killed by a motorized vehicle when crossing a road (n = 1). Another three individuals were lost due to unknown reasons, which may have been due to premature transmitter failure, mortality, or the animal moving beyond radio signal despite extensive search efforts. Thus, we only successfully recaptured and re-implanted five individuals (M01 once, M02 twice, M07 once, M27 once, and M33 twice). At the end of the study, only one individual was successfully recaptured to remove the transmitter.

### Data Collection

We used very high frequency radio-telemetry to locate each telemetered individual on average every 24.20 hours (SE ±0.41, 0.17-410.0 hours; see Supp. Figure 2 for distribution of tracking time lags). We aimed to locate each individual’s shelter locations once each day during the daylight (06:00-18:00 h); however, we were occasionally (n = 34 days) unable to locate a snake for several consecutive days when we were unable to obtain radio signal due to an individual having moved far away or deep underneath a large structure. There were also a few occasions where we were unable to track snakes due to prolonged and heavy rainfall (n = 4 days), as the moisture damages equipment, or other reasons (n = 4 days). We additionally located snakes nocturnally (18:00-06:00 h) ad hoc and in an attempt to observe nocturnal behaviors and movement pathways when animals were active. We defined fixes as any time a telemetered individual was located, and relocations (i.e., moves) as the occasions where we located an individual >5 m from its previous known location.

Each day we manually honed in on signal via a radio receiver to locate individuals (as described by Amelon et al., 2009), and recorded locations in Universal Transverse Mercator (UTM; 47N World Geodetic System 84) coordinate reference system with a handheld global positioning system (GPS) unit (Garmin 64S GPS, Garmin International, Inc., Olathe, Kansas) directly above the sheltered snake. We generally approached within one meter of sheltering snakes during daylight to precisely record shelter locations and identify shelter type. Since we could not visually confirm snake locations, we methodically eliminated all possible locations where the snake could possibly be while at close range with the minimum possible gain on the radio receiver.

Telemetered kraits tended to be inactive and sheltering underground during the daylight, thus we were confident that our diurnal location checks would not affect their movements. However, in some cases we resorted to determining an individual’s location via triangulation, where multiple lines cast from different vantage points towards the snake intersect on the snake’s location on the GPS, allowing us to determine the animal’s coordinate location from approximately 10-30 m away. This helped ensure that we recorded locations with greater accuracy when snakes sheltered underneath large buildings, as it allowed us to move away from large structures that hindered the GPS accuracy. This technique was also implemented during some nocturnal location checks when a snake was believed to be active among dense vegetation, in an attempt to prevent disturbance of the animals’ natural behavior. While we did hope to gain visual observations of active individuals during the night, we exercised more caution during nocturnal location checks, typically maintaining a minimum distance of approximately 5 m in attempt to lessen the chances of disturbing an active individual’s behavior. If the animal was active we recorded the animal’s observed behavioral state (i.e., moving, feeding, or foraging). When the radio signal was stable and the individual was not visible, we recorded the animal’s behavior as “sheltering”. We strived for an accuracy of <5 m GPS accuracy when feasible during each location check.

For every location fix we recorded the time (dd/mm/yyyy hh:mm), location (Universal Transverse Mercator, UTM; World Geodetic System 84), relocation distance (straight-line distance between the last known location to the new location, relocation/move defined as >5 m difference), and land-use type (e.g., mixed deciduous forest, human-settlement, semi-natural area, agriculture, plantation; see Supp. Figures 3-4 for photos of land-use types), behavior (e.g., sheltering, moving, foraging, or feeding), and shelter type (e.g., anthropogenic, burrow, or unknown, note we also recorded if we suspected the shelter to be part of a termite tunnel complex due to a close proximity to a visible termite mound; Supp. Figure 5).

During each location check we recorded the straight-line distance between the current and previous locations (distance moved/step length) with the GPS device. We then used step-lengths to summarize their movements by estimating the mean daily displacement (MDD; the total distance moved divided by the number of days the snake was located) and mean movement distance (MMD; the mean relocation distance, excludes distances ≤5). In order to limit biases due to some snakes being located multiple times within a given day/night, we limited our sample for estimating MMD and MDD to only include a single location per day. This was accomplished by manually removing “extra” nocturnal location checks that occurred within the same day, making sure to have all shelter relocations present within the dataset. When calculating MDD, we used the total number of daily location checks rather than the number of days between the individual’s tracking start and stop date since there were some days where individuals were not tracked. We also used the same one location check per day dataset to calculate movement/relocation probabilities and to examine each individual’s MMD, MDD, and relocation probability for the overall tracking duration as well as for each season.

When feasible, we positioned a Bushnell (Bushnell Corporation, Overland Park, Kansas) time lapse field camera (Trophy Cam HD Essential E3, Model:119837) with infrared night capability on a tripod spaced 2-5 m from occupied shelter sites. We positioned the cameras so that we may gather photos of the focal snake as it exited the shelter site and/or behaviors exhibited near the shelter. We programmed the cameras using a combined setting, including field scan, which continuously captured one photo every minute, along with a motion sensor setting, which took photos upon movement trigger outside of the regular 1- minute intervals.

### Space use and site fidelity

All analyses and most visualizations were done in R v.4.0.5 using RStudio v.1.4.1106. We attempted to estimate home ranges for the telemetered *B. candidus* individuals using autocorrelated kernel density estimates (AKDEs) using R package *ctmm* v.0.6.0 (Calabrese, Fleming & Gurarie, 2016; Fleming & Calabrese, 2017) in order to better understand the spatial requirements of *B. candidus*. However, examination of the variograms produced after running the best performing movement models revealed that the majority of the variograms had not fully stabilized (i.e., limited evidence of range stability in our sample), and many individuals had extremely low effective sample sizes (21.82 ±9.75, range = 1.49– 135.75; Supp. Table 4). Therefore, we do not report home ranges in this text, as the AKDE estimates would violate the assumption of range residency and either underestimate or misrepresent *B. candidus* spatial requirements. We also examined the speed estimates resulting from fitted movement models. Resulting variograms and tentative home range estimates are included in a supplementary file for viewing only (Supp. Figure 6, Supp. Table 4). The original code is from Montaño et al., (2021).

Since our data was not sufficient to estimate home range size for the telemetered *B. candidus*, we instead used Dynamic Brownian Bridge Movement Models (dBBMMs) with the R package *move* v.4.0.6 (Kranstauber, Smolla and Scharf, 2018) to estimate within study occurrence distributions. We caution readers that these are not home range estimates but instead modeling the potential movement pathways animals could have traversed (Kranstauber et al., 2012). Use of dBBMMs not only allows us to estimate occurrence distributions for each individual, thus helping us better understand the animal’s movement pathways and resource use, but it also allows us to examine movement patterns through dBBMM derived motion variance (Silva et al., 2018; Smith et al., 2021). We selected a window size of 19 and margin size of 5, to catch short resting periods with the margin, while the window size of 19 is long enough to get a valid estimate of motion variance when the animals exhibit activity/movement. Contours however are somewhat arbitrary; therefore, we used three different contours levels (90%, 95%, 99%) to estimate dBBMM occurrence distributions (using R packages *adehabitatHR* v.0.4.19, and *rgeos* v.0.5.5), and show the sensitivity to contour choice.

All movement data, either including initial capture locations or beginning with the first location check ∼24 h post release, was used for production of both the AKDEs and dBBMMs for each individual. We also estimated dBBMM occurrence distributions for each telemetered individual with the exception of M29, which only made three small moves within a burrow complex during the short time he was radio-tracked before transmitter failure.

We compared space use estimates to two previously published *B. candidus* tracking datasets (Mohammadi et al., 2014; Knierim et al., 2018), and one unpublished dataset shared on the Zenodo data repository (Smith and Knierim, 2021), originating from the Sakaerat Biosphere Reserve (approximately 41 kilometers to the south of our study site): two adult males from within the forested area of the reserve [one tracked every 27.8 ± 0.99 hours over a period of 103 days, the other tracked every 38.63 ± 11.2 hours over a period of 30.58 days] (Mohammadi et al., 2014; Smith and Knierim, 2021), and a juvenile male from agriculture on a forest boundary [tracked every 50.19 ± hours for 66.91 days] (Knierim et al., 2018). The previous studies on *B. candidus* only tracked the movements of a single individual each, had coarser tracking regimes, and used traditional–fundamentally flawed methods (Silva et al., 2020; Crane et al., 2021)–to estimate space use (Mohammadi et al., 2014; Knierim et al., 2018). Therefore, we ran dBBMMs with these previous datasets using the same window and margin size (ws = 19, ms = 5).

To quantify site reuse and time spent at sites (residency time) we used recursive analysis with the R package *recurse* v.1.1.2 (Bracis et al., 2018). We defined each site as a circular area with a radius of 5 m around each unique location (matching the targeted GPS accuracy). Then we calculated each individual’s overall number of relocations, each individual’s total number of relocations to each site, and each individual’s site revisit frequency and residency time at each unique site. Then we plotted revisited locations on a land-use map with space use estimates (95% and 99% dBBMM) in an attempt to help identify and highlight activity centers for telemetered individuals (see Supp. Figures 7-13).

### Habitat selection

We used Integrated Step Selection Function models (ISSF) to examine the influence of land-use features on the movements of *B. candidus* at both the individual and population levels. We included movement data from all male individuals that used more than one habitat feature in our ISSF analysis. Therefore, we excluded F16 and M29 who both only used settlement habitat. Excluding M29 was justified by the individual having been tracked for the shortest duration (19 days) and had the fewest number of moves (n = 3), thus there were not enough relocations for ISSF models to work effectively. Using modified code from Smith et al. (2021) that used ISSF with Burmese python radio-telemetry data, we used the package *amt* v.0.1.4 (Signer, Fieberg and Avgar, 2019) to run ISSF for each individual, with Euclidean distance to particular land-use features (natural areas, agriculture, settlement, buildings, and roads) to determine association or avoidance of features. Cameron Hodges created all land-use shape files in QGIS by digitizing features from satellite imagery and verified all questionable satellite land-use types via on-ground investigation.

The semi-natural areas, plantations, mixed deciduous forest and water bodies (such as irrigation canals and ponds which have densely vegetated edges) were all combined into a single layer of less-disturbed habitats which we refer to as “natural areas”. All feature raster layers were then converted into layers with a gradient of continuous values of Euclidean distances to the land-use features, and were inverted in order to avoid zero-inflation of distance to feature values and to make the resulting model directional effects easier to more intuitive. We were able to generate 200 random steps per each observed step (following Smith et al., 2021), due to the coarse temporal resolution of manually collected radio-telemetry data (i.e., we were not computational limited when deciding the number of random locations). Higher numbers of random steps are preferable as they can aid in detecting smaller effects and rarer landscape features (Thurfjell, Ciuti and Boyce, 2014).

To investigate individual selection, we created nine different models testing for association to habitat features, with one being a null model which solely incorporated step-length and turning angle to predict movement (Forester, Im and Rathouz, 2009), five examining land-use features individually (agriculture, buildings, settlement, natural areas, roads), and the other three being multi-factor models. Each model considers distance to a land-use variable, step-length, and turn-angle as an aspect of the model. After running each of the nine models for each individual, we then examined the AIC for each model, point estimates (with lower and upper confidence intervals), and p-values in order to identify the best models for each individual and determine the strongest relationships and trends among the samples. We considered models with Δ AIC <2 as top performing models.

We then investigated habitat selection at the population level, including all radio-telemetered except for M29 and F16. We used code from Smith et al. (2021), which was modified code originally from Muff et al. (2020), using a mixed conditional Poisson regression model with stratum specific effects. This model was essentially the equivalent to an integrated step selection model at the population level. Both the step (strata), and the individual (individual ID) are modelling using Gaussian processes. As used for the individual level ISSF, we generated 200 random steps from each location, with a Gamma distribution for step length and Von Mises distribution for turn angle. We created five single factor models using the same land-use features used with the individual level ISSF (i.e., agriculture, buildings, natural areas, roads, and settlements [via the same inverted distance to land-use feature rasters]) with individual random intercepts and slopes. Following Muff et al., (2020), we set a fixed prior precision of 0.0001 for the stratum-specific random effect (i.e., step). We used a Penalized Complexity prior, PC (1, 0.05), for the other random slopes (i.e., individual), and uninformative normal priors, Normal (0, 103), for the fixed effects, as was done by Smith et al., (2021). We used integrated nested Laplace approximations with the INLA package v.20.03.1748 (Rue, Martino & Chopin, 2009) to fit all the models.

### Seasonality

We classified three four-month seasons: wet season (01 June – 01 October), cold season (01 October – 01 February), and hot season (01 February – 01 June). Although we acknowledge that Mean Movement Distance (MDD) and Mean Daily Displacement (MDD) are highly sensitive to tracking regime and tracking duration (Secor, 1994; Rowcliffe et al., 2012), we use these metrics as we attempt to maintain standard daily tracks throughout the study period. We calculated MMD and MDD for each individual within the defined seasons, using the methodology described for gaining overall MMD and MDD, and movement probabilities for each individual during each season (the total number of fixes divided by the number of “relocations”), in order to examine the raw data for possible movement trends.

We also examined dBBMM derived motion variance of all individuals to examine potential trends in temporal activity, and documented observations which may help determine when breeding takes place (e.g., conspecific interactions, presence of sperm plugs in males).

## Results

### Movement summaries

We tracked 14 individuals for an average of 105.56 SE ±15.16 (range = 19-218) days. During the tracking period, we located individuals on average every 24.03 ±0.43 hours, and detected 25.71 ±4.09 moves per individual with a mean step-length of 24.96 ±2.05 m.

We gathered a total of 1,505 fixes and 324 relocations (>5 m, counting initial capture locations), with 1,381 fixes during the daylight and 124 fixes during the night; 752 fixes occurred during the cold season, 445 during the hot season, and 308 during the wet season.

Males (n=13) moved an average of 2,772.31 ±443.88 m during their tracking durations, and had a MMD of 117.78 ± 8.23 m (range = 6–1130 m), a MDD of 27.48 ±2.36 m, and mean daily movement probability of 0.23. Adult males tended to exhibit higher mean motion variance and MMD than juvenile males (Table 2). Mean motion variance was highest in M32 (9.63 ±2.34 m), a large adult male which also had the greatest MMD (259.65 ±57.67 m) and MDD (60.99 ±17.37 m). Speed estimates were on average 21.82 ±9.75 m/day (1.49–135.75), but were incalculable for eight individuals (due to lack of model fit). Compared to MDD and MMD, the existing speed estimates (n = 5) appear very weakly connected to MMD (Supp. Figure 14).

Mean motion variance for all telemetered *B. candidus* was low, at 1.70 ±0.18 m (5.52 x 10^-5^–89.73 m). Mean motion variance was lowest for the single telemetered female (mean 0.16 ±0.04 m), who remained within the same shelter complex (under the F1 building near the northern entrance) for the majority of her tracking duration, including 85 consecutive days (25 January – 20 April 2019) spent there.

### Occurrence distributions and site fidelity

Individual dBBMM occurrence distributions varied greatly, with the smallest (excluding M29) being the female (F16) with a 99% confidence area of 0.42 ha, and the greatest being the 99% confidence area for M32, at 119.55 ha (Figure 2). We removed one male (M29) from this summary because the tracking duration was only 21 days with a single small relocation prior to returning back to the previous site. The telemetered male *B. candidus* (n = 12) had a mean 90% dBBMM confidence area of 6.66 ha (±2.41, 1.06- 29.81), 95% of 11.22 ha (±4.37, 1.52-56.20), and a 99% of 22.33 ha (±9.21, 2.52-119.55; Table 1).

**Figure 2.**
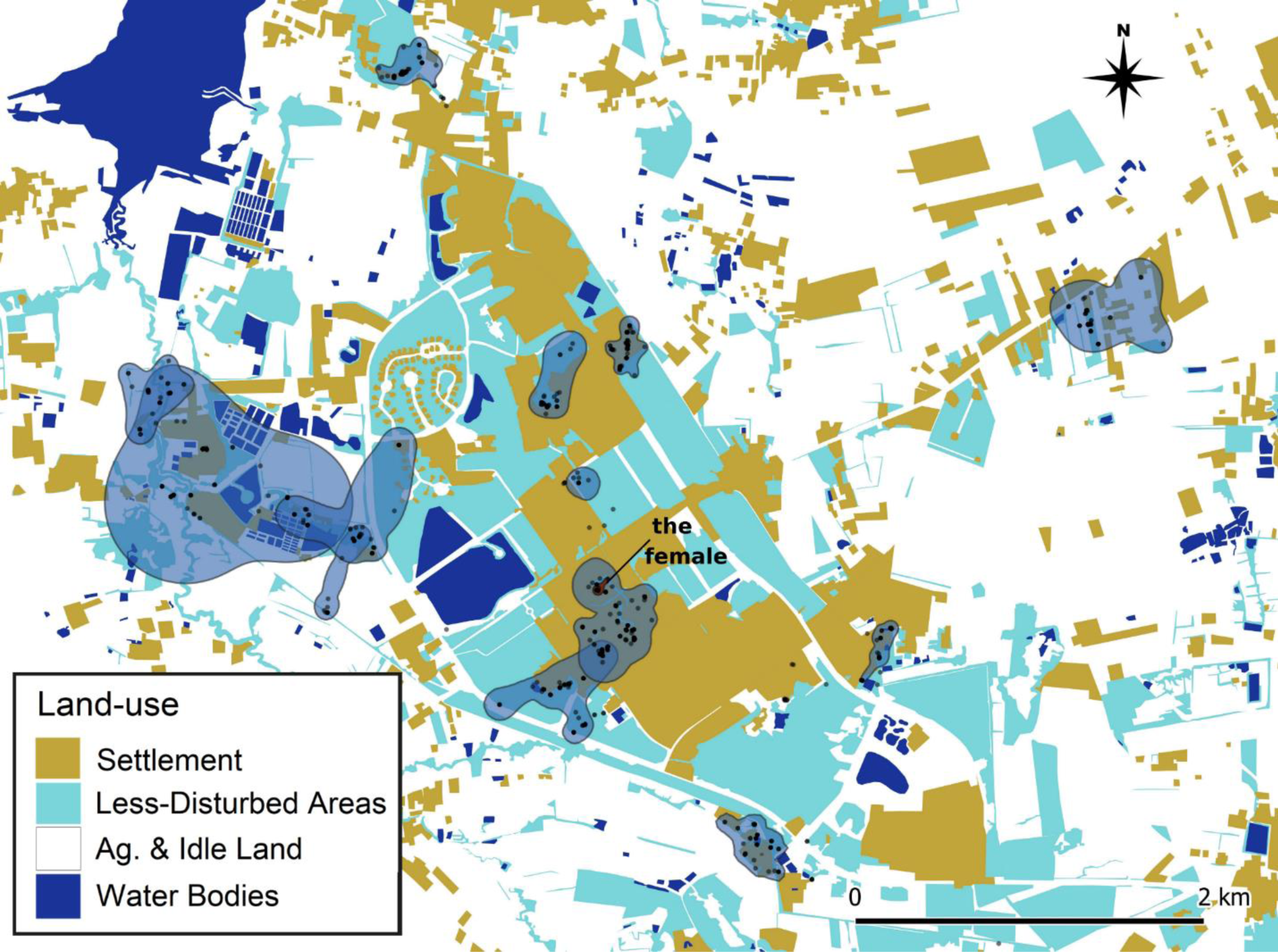
dBBMM occurrence distributions (male 99% confidence area polygons in blue, female 99% confidence area polygon in red) and location fixes (semi-transparent black dots) for each radio-tracked *B. candidus* individual in Muang Nakhon Ratchasima, Thailand.

Compared to our male occurrence distributions (95% confidence area mean = 11.22 ±4.37 ha, range = 1.52-56.2 ha), the occurrence distributions produced for the male *B. candidus* living in less-developed environments at the Sakaerat Biosphere Reserve (n = 3, 95% contour estimate mean = 8.49 ±2.4 ha, range = 4.72-12.95 ha) were very similar.

All except one (M36) of the 14 radio-tracked individuals revisited at least one shelter site during tracking. For these individuals (excluding M29), the overall mean number of site revisits was 18.67 (range = 2-46), with an overall mean site revisit frequency of 15.45 ±3.87 days (1.18–50.57 days). Mean average time telemetered *B. candidus* remained within a shelter was 7.69 ±1.98 days (1.75–30.475).

### Habitat use

Habitat use varied across individuals; however, the most frequently used habitat type overall was human settlement, with 51.2% of all fixes (Figure 3A). Semi-natural areas were the second most commonly used habitat (25.2%), closely followed by mixed deciduous forests (22.8%). The least used land-use types were agriculture (0.5%) and plantation forests (0.3%). Of the points among human settlement habitat, 558 (72.47%) were associated with buildings and 99 (12.86%) were associated with concrete drainage ditches, sidewalks, or other concrete structures.

**Figure 3.**
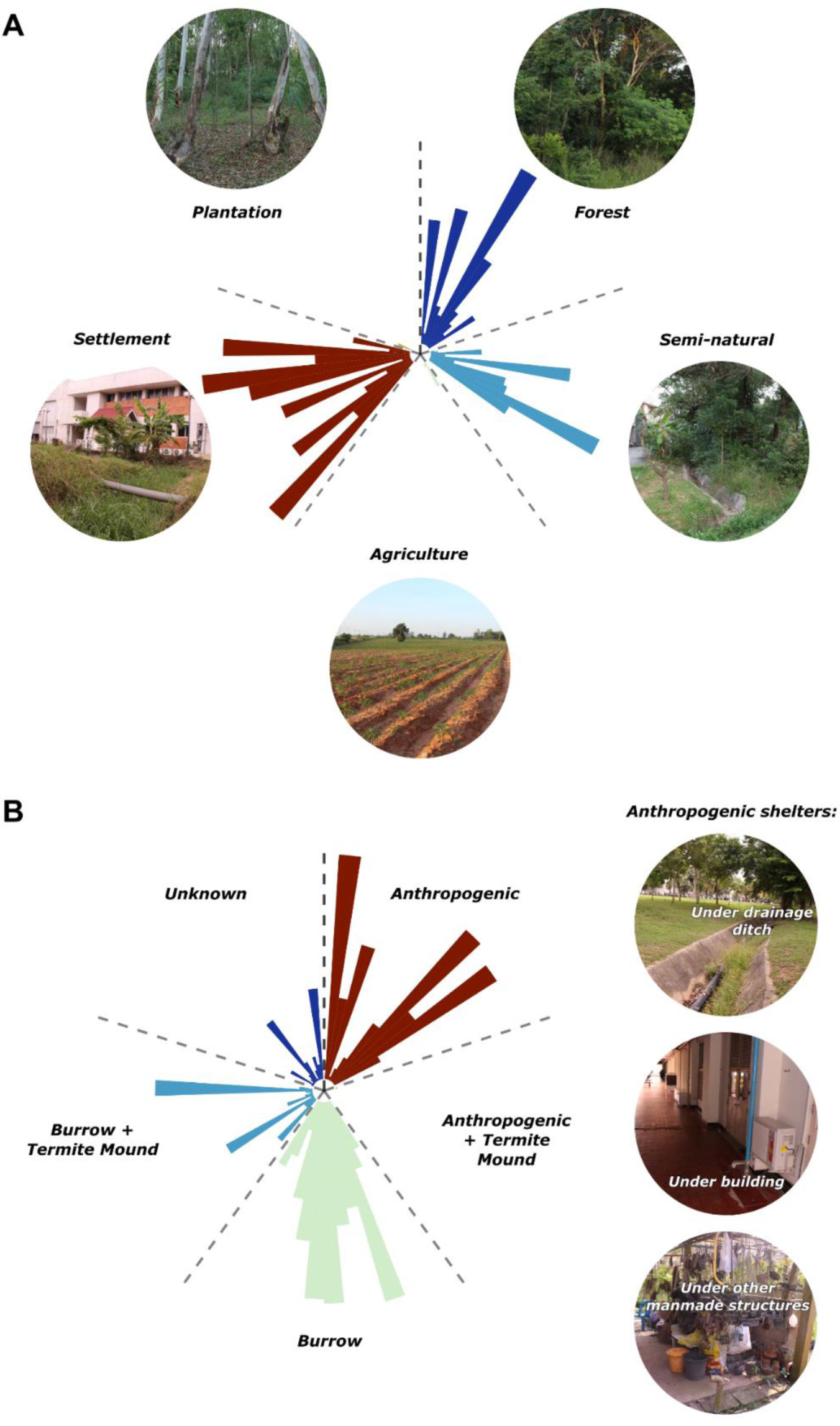
Proportional **A)** habitat use and **B)** shelter site use proportions for each telemetered *B. candidus* individual from Muang Nakhon Ratchasima, Thailand.

### Shelter use

We determined individuals to be sheltering during 1,443 fixes. The most commonly used shelter type, with 582 (40.3%) fixes, were burrows, however, use of anthropogenic shelters was nearly equal, with 559 (38.7%) fixes (Figure 3B, Supp. Figure 5, Supp. Table 6). Anthropogenic shelters included fixes where snakes were directly underneath buildings (507), concrete drainage ditches (37), sidewalks (2), or other anthropogenic structures, while burrows included burrow systems and tunnels excavated by animals, such as rodents, but did not include shelters which appeared to be part of termite mound tunnel systems.

Termite mounds/tunnels made up 217 (15%) of the shelter fixes. We were unable to identify shelter types during 85 (5.9%) of the fixes (though in as much as 55% of these “unknown” shelter fixes the snake was suspected to be sheltering among dense vegetation).

One-hundred and fourteen of the shelters classed as non-“anthropogenic” were within a short distance (<5 m) of concrete structures, such as drainage ditches or buildings, with 37 of the “burrow” and two “unknown” shelters within a single meter of a building, nine in or within a meter of a concrete drainage ditch, and 19 within a meter of a paved sidewalk.

### Trends in foraging sites

Since we attempted to limit nocturnal location checks, it was uncommon for us to track an individual when it was active. Of the 38 occasions where we observed telemetered snakes moving, foraging, feeding, or otherwise active, nine occurred within a single meter of a paved sidewalk, 10 were associated with buildings (either inside or < 1 m to a building edge), and eight occurred in or within a single meter of a concrete drainage ditch (note these instances include initial capture locations). In total, 22 of 37 observations were associated (< 1 m) with concrete structures of some kind. We also gained five observations of *B. candidus* foraging or moving within agriculture (two within cassava fields, one within a fallow field, one among a fishery, and one on a road-side among a grass field). Several of the other observations occurred near the edge of a body of water, and another two telemetered individuals were observed moving within a meter of a chicken coop.

We recorded two individuals feeding on snakes during the study; both were within anthropogenic land use types (fully reported elsewhere by Hodges, D’souza and Jintapirom, 2020; Hodges et al., 2021). The first occurred in a concrete gutter within 2 meters of student housing. The second event occurred within an open atrium within a faculty office building.

### Habitat selection

All twelve of the *B. candidus* individuals included in the ISSF analysis exhibited positive association with natural habitats, and all but two of the individuals (M27 and M35) showed positive association with anthropogenic structures (Supp. Figure 16). Five different models best explained habitat selection across individuals (Supp. Table 7). Top models included model4 (buildings), model6 (natural), model7 (agriculture, natural, and buildings), model8 (roads, buildings, and natural), and model9 (roads, agriculture, and natural). Model8 was the top model for five of the individuals (M02, M14, M27, M28, and M36) and showing a positive association for each feature included in the model (roads, buildings, and natural), with the exception of M02 and M28 which showed negative association with roads and M27 which showed negative association with buildings. Model7 was the top model for four individuals (M01, M07, M12, and M22) with a positive association with buildings and natural features but a negative association with agriculture. Model9 (roads, agriculture, and natural) was the best model for M35 (showing association with each feature). Model6, which was a single factor model examining the interactions with the distance to natural habitat features, was the best model for M32 (showing negative association), while model4, a single factor model using distance to buildings, was the top model for M33 (revealing positive association).

Credible intervals were quite broad and sometimes overlapped zero for several of the individual’s models, thus limiting our ability to draw inferences. Much of the model uncertainty is likely resulting from the coarse resolution tracking data and the few and infrequent movements by the study animals. Interactions between step-length and distance to land-use features appeared nonexistent. However, the population level model revealed a potentially broader trend while absorbing some of the individual heterogeneity (via the random effect), with weak but positive associations with natural habitats (mean estimate = 0.0235, 95% CrI 0.0114-0.0416), buildings (mean estimate = 0.0094, 95% CrI 0.0038-0.0165), and settlements (mean estimate = 0.0059, 95% CrI 0.0001-0.0125). A possible weak negative association with agriculture was also present (mean estimate = -0.0018, 95% CrI -0.0058-0.0020; Figure 4). Estimates for the influence of distance to land-use features on step lengths resulting from the population level model were still ambiguous, as all estimates were near zero and had confidence intervals overlapping zero.

**Figure 4.**
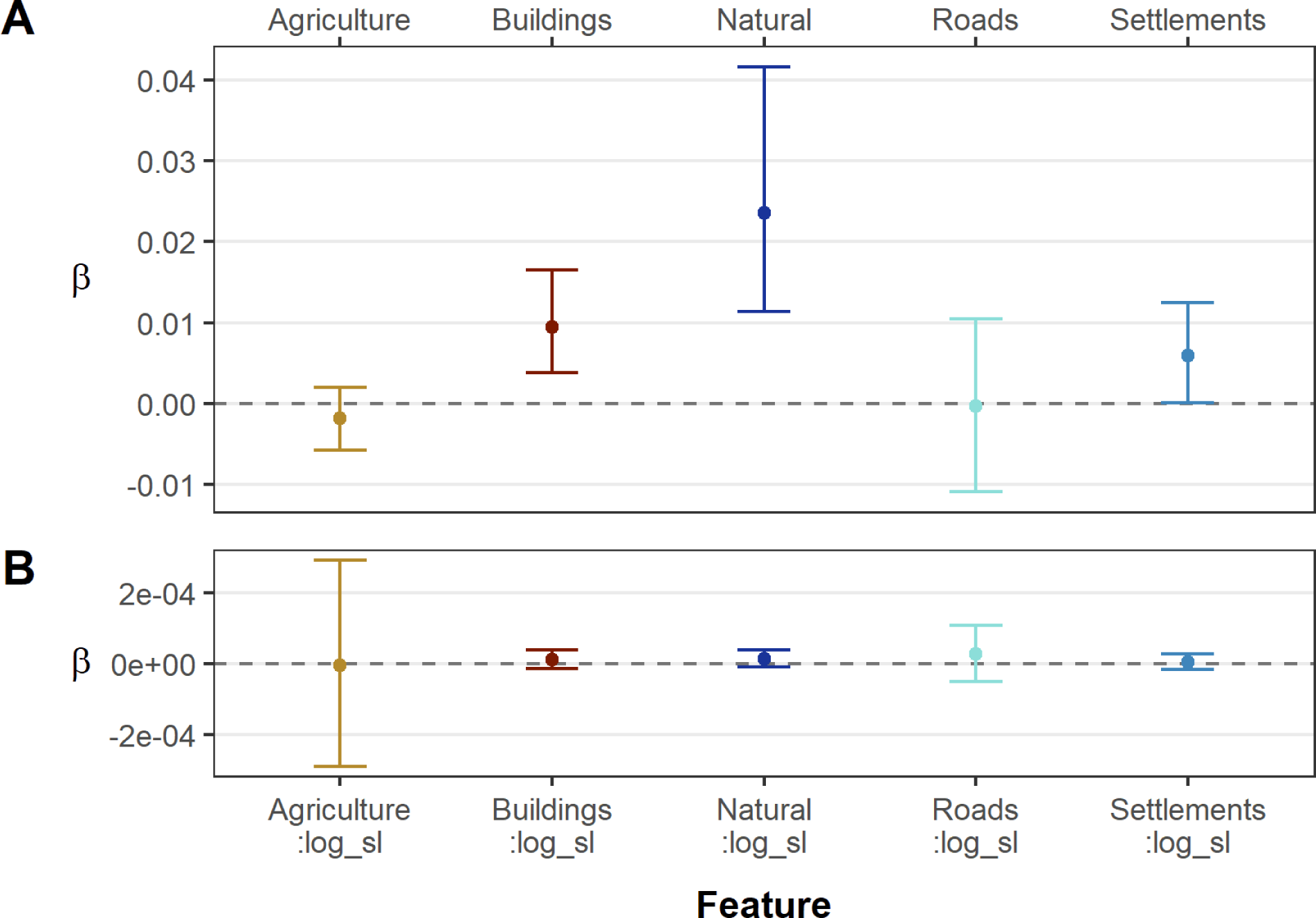
Population level ISSF model results based on distance to habitat features. **A)** Point estimates and 95% confidence intervals for habitat selection. Positive estimates suggest association with the habitat feature. **B)** Estimates for the influence of distance to habitat features on step lengths.

### Seasonality and Temporal Activity Patterns

Males moved furthest distances (MMD) in the cold season, but longer distances per day (MDD) and more frequently in the wet season. In contrast males moved least (MMD and MDD) in the hot season (Figure 5B, Supp. Table 8). Males were more likely to move between any given daily datapoint in the wet season than in other seasons.

**Figure 5.**
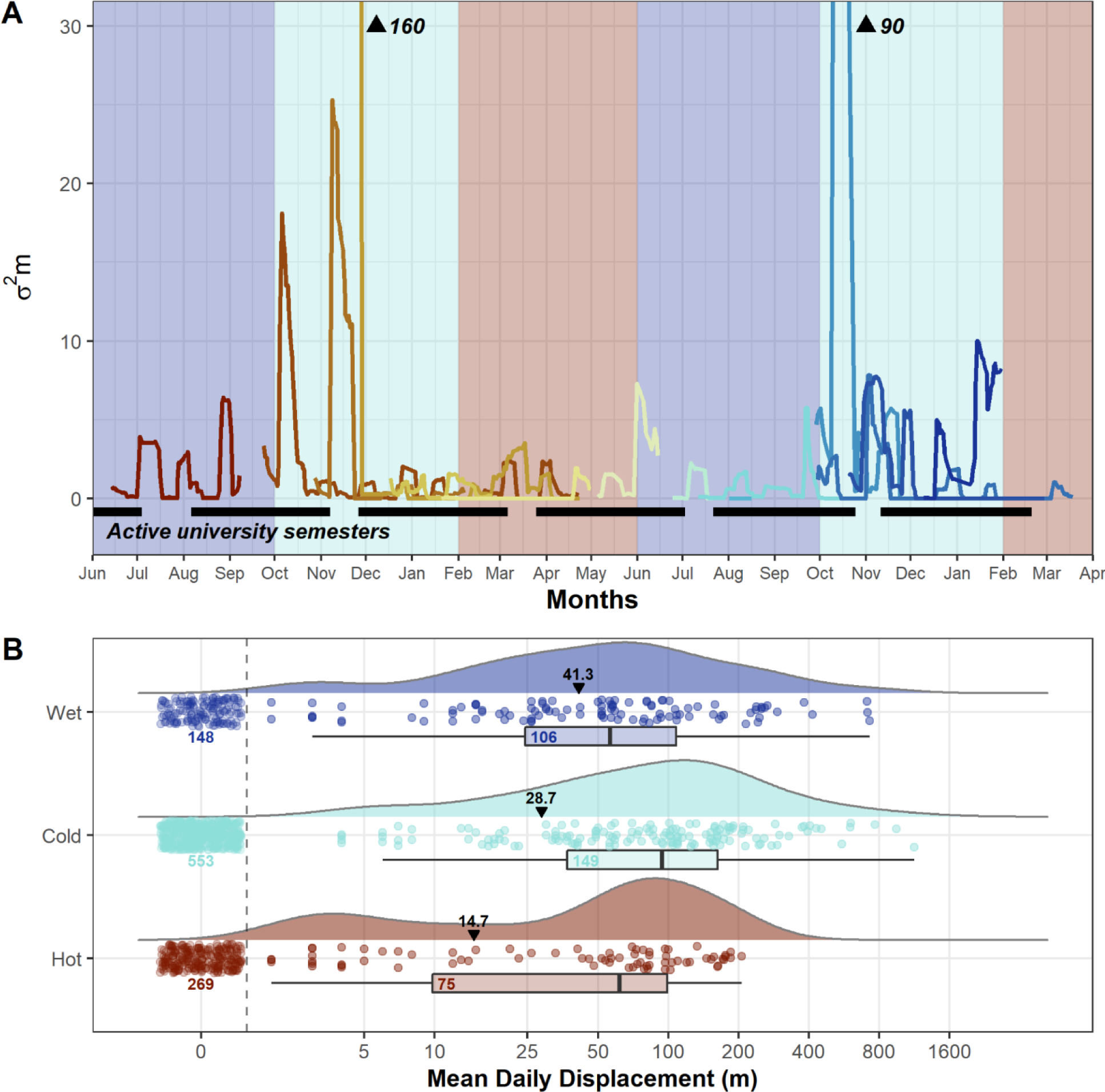
**A)** Motion variance for each individual throughout the study period, with bars indicating when university semesters were in session and background colors corresponding to season: blue = wet, light blue = cold, red = hot. Numeric annotations indicate two peaks in motion variance for, M12 and M32 respectively, that exceed the limits of the plot. **B)** Raincloud plots showing the Mean Daily Displacement in each season for the 13 tracked males. Box and density plots are plotted excluding the non-moves, and labels describe the number of points in each season. Non-moves are displayed as jittered points to the left of the dashed line, with the count displayed below. Black labels show the overall mean per season, including all moves and non-moves.

Mean motion variance was highest during the cold season (2.579 ±0.332 m), where four larger adult males peaked in motion variance resulting from particularly large movements coinciding within the first two months of the cold season (Figure 5A). There notably were several particularly high peaks in motion variance from a few different individuals (M02, M07, M12, and M32) both years. These four highest peaks all coincide within the first two months of the cold season, October and November. Similarly, twelve of fourteen particularly large movements (≥395 m, by 6 individuals) documented occurred within the late wet season (n = 4) or early cold season (n = 8), with the remaining two occasions occurring within the late cold season (January). In contrast, motion variance was lowest in the hot season (0.481 ± 0.042 m), and the wet season average was roughly halfway between the other seasons (1.174 ± 0.105 m).

In contrast to large moves, six of the male Malayan kraits showed prolonged stationary periods, remaining inactive within the same shelter for ≥20 consecutive days (mean = 35.05 ±7.76 days, range = 20-94 days, n = 9). The prolonged stationary periods occurred eight times in the cold season, (four of those eight occurring late cold season just prior to the hot season), and once in the hot season.

### Shelter Emergence Times

We gathered a total of 1,160,970 time-lapse camera photos. Of these, we could only identify focal animals in 75 photos, from six different individuals, on 14 different occasions (i.e., independent nights), with a mean of 5.36 (range = 1-18) photos each occasion. During different occasions, individuals generally either peaked their heads out and slowly exited shelter sites (n = 7), or simply exited the shelter site and immediately moved away (n = 4), not to return again. Two individuals (M28 and M36) were photographed active during the night, presumably foraging near to the shelter site, before returning to the same shelter. On another occasion an individual (M12) spent a few minutes lying just outside the shelter before moving off.

On camera, individuals tended to exit shelters and begin moving and/or foraging shortly after sundown, with all photographed individuals moving outside the shelter sites between 19:00 and 22:30 h, and with shelter site activity peaking at 19:30 h (Figure 6).

**Figure 6.**
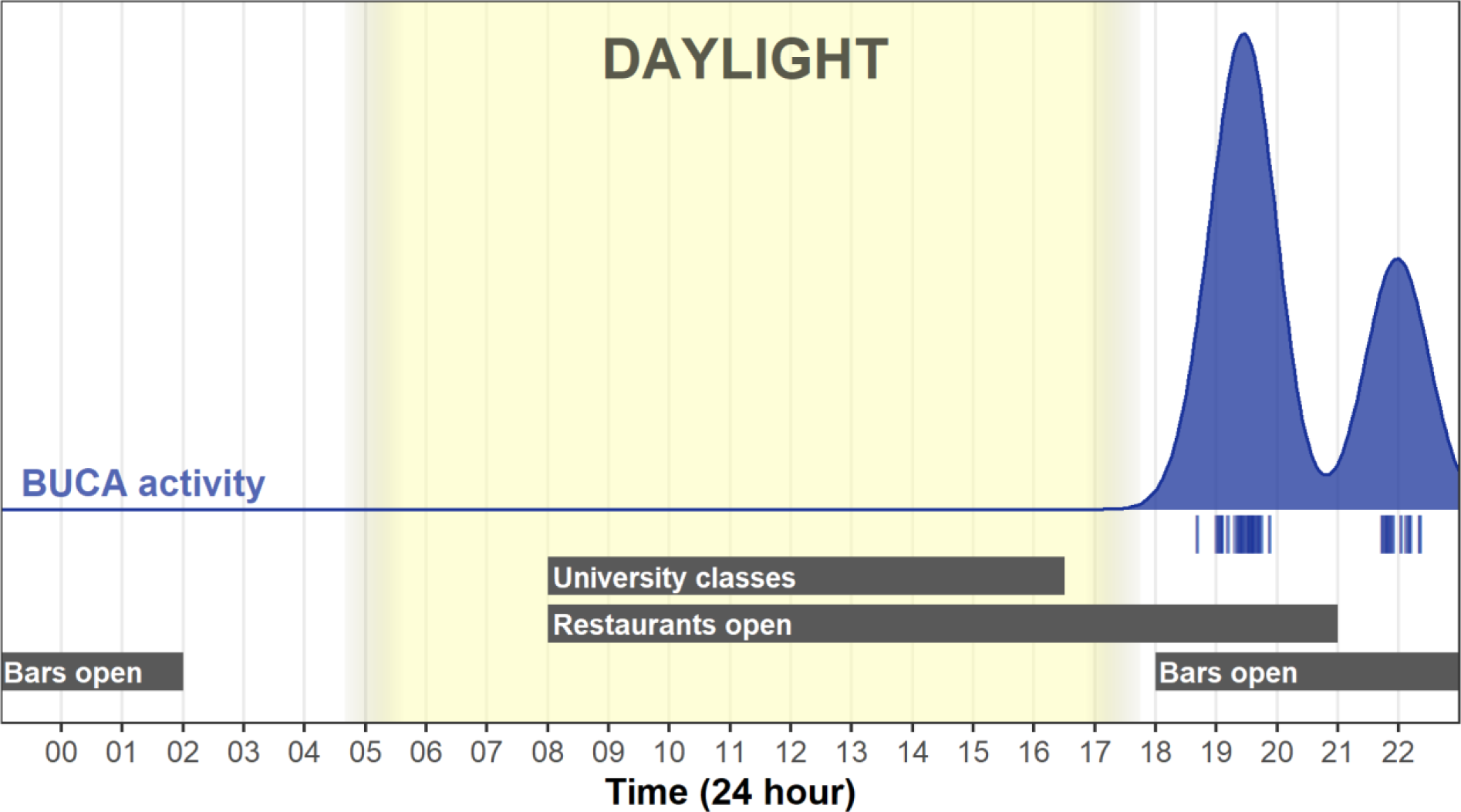
Daily shelter site emergence activity patterns of *B. candidus* based on observation via camera traps (blue density plot and points). Below are several potential drivers of human movement about campus, as well as the daylight hours (the gradient shows the variation in sunrise/set during the year).

## Discussion

This study yields insight into a secretive snake species of medical importance –*Bungarus candidus*. We use a variety of approaches (i.e., dBBMMs, ISSFs, and AKDEs) to analyze low-resolution animal positional data. Despite identifying site fidelity, we were unable to estimate home ranges (via AKDE) of *B. candidus* due to a surprising lack of evidence of range stability, likely due to limited number of relocations, resolution, and tracking duration. However, dBBMM movement pathway estimates suggest an average area of 11.22 ±4.37 ha (95% confidence area) used during the study period per male individual. *Bungarus candidus* in our study were seasonal and frequently sheltered, foraged, and remained in human settlements, with high site fidelity and long-term use of shelters in areas constantly co-habited by humans. Despite *B. candidus* consistently occupying areas near humans (and potentially being most active during times of relatively high human traffic/movement) there were no incidents of snakebite attributable to *B. candidus* in our site during the study period.

### Space use

Space use is frequently dependent on the environment (Van Moorter et al., 2016); however, we witnessed very little difference between our kraits’ confidence areas (n = 13, 95% confidence area mean = 11.22 ±4.37 ha), and the previous studies in much less urban areas (n = 3, 95% contour estimate mean = 8.49 ±2.4 ha). This similarity appears to contrast with findings from Tucker et al. (2018), who showed that mammals move less in anthropogenic environments. While the comparison is limited by the sample size from less-developed areas and differences in tracking duration, the initial lack of difference may indicate krait movement patterns are insensitive to human presence, or that changes are occurring on a scale undetectable by our dataset (e.g., movement changes occurring in <24 h, changes in activity times). A lack of flexibility may lead to heightened vulnerability to human-snake conflict (Fahrig, 2007), and also presents a foundation for exploring how such conflict may be predicted by human behaviors.

### Site Fidelity and Habitat Selection

All but one of the 14 radio-tracked individuals exhibited site fidelity. Snakes revisited both natural and developed areas repeatedly, specifically reusing termite mounds, tunnel systems, and crevices under anthropogenic structures (Figure 3B, Supp. Figure 5).

We found most individuals to be associated with natural areas, commonly moving through and sheltering among available mixed deciduous forest and semi-natural areas. But, most individuals also tended to use settlements more so than less-disturbed habitats, and most individuals were positively associated with anthropogenic structures, such as houses and university buildings. Snakes commonly used these areas as shelter sites (likely leading to the building association in the population model). Several studies document habitat use and movement with snakes in landscapes associated with humans (Butler, Malone and Clemann, 2005; Anguiano and Diffendorfer, 2015), though relatively few evaluated or even documented sheltering under concrete anthropogenic structures (Lee, Park and Sung, 2012; Gerke, Hinton and Beasley, 2021). In cooler temperate areas, some snakes frequently used anthropogenic structures as refugia, as they provided more suitable thermal qualities than the available natural shelters (Keller & Heske, 2000; Lelièvre et al., 2010). More relevant to our study, Wolfe et al, (2018) reported that several telemetered *Pseudonaja affinis*, a large diurnal elapid species from Australia capable of surviving in urban areas, occasionally sheltered underneath housing and paving stones. Our study demonstrates that even in a warm tropical climate, potentially dangerous snakes may be attracted to buildings and may use them as shelters even more frequently than available natural refuges.

Why are kraits using human settlements? These sites may modulate temperature by increasing ambient surrounding landscape temperature (Scott et al., 2017). So perhaps Malayan kraits are using these insulated concrete structures to thermoregulate. While some suggest thermoregulation is less important to reptiles in tropical environments (Slip and Shine, 1988; Shine and Madsen, 1996), the concrete structures may aid animals in avoiding heat (Luiselli and Akani, 2002). Findings from several studies suggest that habitat and shelter selection by some snakes is influenced by prey availability (Whitaker and Shine, 2003; Heard, Black and Robertson, 2004), though in temperate areas it appears that thermoregulation needs may outweigh the benefits from selecting habitats based on prey availability (Blouin-Demers and Weatherhead, 2001; Sperry and Weatherhead, 2009). Concrete structures likely house prey species such as other snakes, anurans, lizards, and mammals. Rats for example often use these structures because they fulfill three criteria: access to food via refuse, access to water via runoff, and access to shelter as a central foraging location (Oca et al., 2017). Thorough investigations into these dynamics have yet to be held in the tropics, where thermoregulation, though still important (Luiselli and Akani, 2002; Anderson et al., 2005), likely has less of an impact on snake habitat use. Our study does provide evidence that *B. candidus* do forage in areas proximal to shelter sites, sometimes even returning to the same shelter site following active foraging activities (as observed through several camera trap images, also see Hodges et al., 2021).

We found no clear evidence for road avoidance and limited evidence of attraction to roads in some individuals –but not the population model. This may help explain the numerous documented *B. candidus* road mortalities in our study site (C.W. Hodges, personal communication, August 2021), though we suspect culverts were used for at least some road crossings, as appears to be the case for *Ophiophagus hannah* (Jones et al., 2021). Furthermore, while our ISSF models did not reveal clear evidence of avoidance of agriculture, despite agriculture being the most widespread land use type in the area, few individuals used it for shelter. Agricultural areas could have occasionally been used as foraging sites during the night (such as observed in M27 and M33), but not as shelter sites due to the lack of available shelters. More commonly, *B. candidus* sheltered in unmanicured field margins, similar to findings from Knierim and colleagues, which observed this in a single telemetered *B. candidus* (Knierim et al., 2018) as well as several radio-tracked *B. fasciatus* (Knierim et al., 2019) living among agriculture. They attributed the non-use of monoculture plots to insufficient shelter availability resulting from frequent and regular disturbance to top soil by human activities. It is also possible that *B. candidus* may have tended to avoid agricultural areas due to the increased risk of mortality (e.g., Crane et al., 2016, Knierim et al., 2017; Hodges, unpublished data) but we lack direct evidence to support this.

### Seasonality

Tracked male *B. candidus* had more frequent but shorter relocations in the wet season, and moved less frequently, but covered greater distances, in the cold season, with the least movement in the hot season.

The wet season may present abundant resources, influencing movement patterns (Wasko and Sasa, 2012; Doherty and Driscoll, 2018). Prey availability can influence snake movement and activity in the temperate (Sperry, Ward and Weatherhead, 2013) and the tropics (Christian et al., 2007; D’souza et al., 2021).

The least resources likely exist in the hot season, coinciding with the least *B. candidus* activity. Other reptiles in northeast Thailand, including *Indotestudo elongata*, (Ward et al., 2021), *Ophiophagus hannah* (Marshall et al., 2020), *Trimeresurus macrops* (Strine et al., 2018), and *Python bivittatus* (Smith et al., 2021) responded similarly to dry seasons. Many reptiles reduce activity during particularly hot and dry periods to conserve energy and water (Christian and Green, 1994; Christian et al., 1995, 2007; Peterson, 1996; Loehr, 2012). *Bungarus candidus* are capable of prolonged inactivity; as we documented six adult males which remained within a single refugia for extended periods (up to 94 days, and this could have continued even longer, as this individual, M35, was recaptured during this period). Snakes may have occasionally exited shelters for nocturnal foraging and returned to the same shelter, but given camera traps failed to reveal movement from shelters, we consider it unlikely (though other exits could exist).

Snakes sheltering long-term may have also foraged and fed fossorially/opportunistically from within the shelter complexes. The long-term dormancy might alternatively be extended mating behavior, though we did not detect conspecifics present during recaptures or shelter site camera trapping.

Peaks in cold season motion variance could be attributed to male *B. candidus* searching for mates over great distances (M02 expelled a sperm plug during processing on 17 September 2018, just prior to cold season). These exceptionally large movements (max of 1,130 m) additionally increased MMD within the cold season. We suspect –but did not confirm– that the telemetered female nested beneath a building foundation in a refuge system where she remained for 85 consecutive days (25 January – April 20 2019). In Thailand, *B. candidus* tend to nest between February and March (Chanhome et al., 2011), and the behavior paralleled nesting *B. fasciatus* documented by Knierim et al. (2019). If females do nest under buildings, there are clear human safety implications, as neonates may enter homes upon hatching between April and May.

The temporal activity patterns presumably impact krait-human conflicts. People should be likely to encounter *B. candidus* in our study site during the wet season, and least likely during the hot season. Tongpoo et al. (2018), showed similar risk trends via hospital records for *Bungarus spp*. (68 of 78 examined bites were by *B. candidus*) bites, with the majority (48.7%) of the bites from kraits occurring in wet season, and the fewest (20.5%) in the hot season.

### Implications for human safety

There is extreme spatial and temporal overlap between humans and kraits in our study, similar to Glaudas (2021) who documented overlap of *Daboia russelii*, another tropical Asian medically significant species. Our findings suggest an increased need for education and awareness among Thai communities, especially when paired with insights provided by Hodges et al., (2021) demonstrating that short-distance translocation for *B. candidus* is ineffective for preventing long-term conflicts. Using these data combined with occurrence data may aid in predicting where snakebites are most likely to occur and perhaps elucidate further preventative methods (Bravo-Vega et al., 2019). Such need for increased data is particularly apparent for other *Bungarus spp*. also known to occupy human-modified areas; for example, *B. caeruleus* that is responsible for thousands of deaths across South Asia annually (Saluba Bawaskar and Himmatrao Bawaskar, 2004; Ariaratnam et al., 2008; Suraweera et al., 2020).

Continual use of buildings for foraging and shelter highlights the need for access to antivenom and precautionary measures to prevent snakebite. In Nepal, Samuel et al., (2020) described how an education program –highlighting flashlight use, wearing boots when working in fields, eliminating openings in housing– led to reduced snakebites and mitigated the consequences. Another study from Nepal identified using a bed net whilst sleeping offered strong protection against snake bite regardless of socio-economic status (Chappuis et al., 2007). As tracked *B*. *candidus* tended to avoid open land features like agriculture, land managers could potentially limit conflict by reducing vegetation cover near buildings–but this is yet untested, and cannot come at the expense of broader biodiversity benefits of university campuses (Liu et al., 2021).

## Limitations

Although the study sample of *B. candidus* (n = 14 [adult males = 9, juvenile males = 4, adult female = 1]) somewhat lower than the average sample for snake spatial studies over the last 20 years; our tracking intensity was comparable to the average (Crane et al., 2021), and we were likely more consistent than average snake telemetry studies.

We attempted appropriate methods for comparing movements across seasons (Noonan et al., 2019), but the models failed as a result of our data structure forcing us to resort to weaker proxies (MMD and MDD). We suspect the short tracking duration (mean = 105.56 ±15.16 days), coarse tracking resolution (mean = 24.03 ±0.43 hours), limited relocations (25.71 ±4.09) and unequal sampling across seasons (308 fixes during the wet season / 1,505 fixes total) led to low effective sample sizes and thus model failure. Our inferences are further limited to shelter site selection as opposed to foraging and finer scale movements.

Prompted by the STRANGE framework (Webster and Rutz, 2020) we consider the following as potentially impacting the generalizability of our findings. **Social background** or **rearing history** likely had no impact on the study with limited evidence of sociality in kraits, and all study animals were free-ranging with limited periods in captivity (mean = 5.02 ±0.61 days). **Trappability** likely influenced the individuals we captured as we could not implement systematic trapping. For example, we more likely tracked individuals nearer humans, evidenced by 10/14 individuals originated from notifications from human-snake interactions. Our results cannot be generalized to female *B. candidus* because we tracked a single tracked female. We doubt that individuals **acclimated or habituated** to observer presence, as individuals were exposed to near continuous presence of human disturbance at the study site. In addition, our focus on day time tracks limited our presence during krait’s activity periods as evidenced by our camera trap data. It is difficult to gauge the impacts of **natural changes in responsiveness** caused by unmeasured biological cycles (e.g., diel cycles, stress response attributed to mating season, prey populations). **Genetic make-up** may have influenced observed behaviors because of the small spatial scale, and high likelihood of closely related individuals. We suspect that **experience** (i.e., learning to avoid humans) played a minor role due to the short duration of the tracking, but radio-transmitter replantation for five individuals may have constituted traumatic events to prompt greater human avoidance; however, this was not apparent in individual step-selection models.

## Conclusion

We identified that tracked krait space use during the study consistently overlapped with human structures and that snakes avoided using open landscape features. We also highlight heightened activity during the wet season and during the early hours of the night, suggesting two peaks in possible human encounters and resulting conflict. But, this marked co-occurrence with humans in the absence of recorded bites suggests an alternative. Redoubling efforts to raise awareness of the habits and likely locations of these medically significant snakes are likely important to maintaining this harmonious balance of human-snake coexistence. Further, investigations into finer scale movements (e.g., building entry routes) could yield insights into further drivers enabling the coexistence we observed.

## Acknowledgements

This project was funded by the King Cobra Conservancy. We would like to thank Suranaree University of Technology (SUT) Institute of Science, Institute of Research and Development, SUT Grounds and Buildings, the Nakhon Ratchasima Zoo, SUT Security, SUT Volunteering, and the Herpetofauna Foundation for their support. We would also like to thank the National Research Council of Thailand for providing the permissions for us to conduct research in Thailand (permit Nos. 0002/27 and 0402/4367).

We specifically thank Dr. Wirongrong Changphet of the Nakhon Ratchasima Zoo for conducting all surgical implantation procedures for this study, Anji D’Souza, Sira Jintapirom, and Porramin Patungtaro for assisting with radio-telemetry, Tyler K. Knierim for providing camera traps, and Inês Silva and Matthew Crane for their help in comprehending home range model results.

## Data Availability

All data, code, and supplementary materials used in this manuscript are provided on the Zenodo data repository (https://doi.org/10.5281/zenodo.5495840).

## Author Contributions

Conceptualization: C.W.H., C.T.S., and B.M.M.; Methodology: C.W.H., C.T.S. and B.M.M.; Data Collection: C.W.H.; Formal Data Analysis: C.W.H. and B.M.M..; Resources: C.W.H.; Writing – Original Draft: C.W.H., C.T.S.; Writing – Review and Editing: C.W.H., C.T.S., B.M.M., and J.G.H.; Visualization: C.W.H. and B.M.M.; Supervision: C.T.S. and J.G.H.; Funding Acquisition: C.W.H. and C.T.S. All authors read and approved the final manuscript.

## Supplemental Figures and Tables

**Supp. Figure 1.**
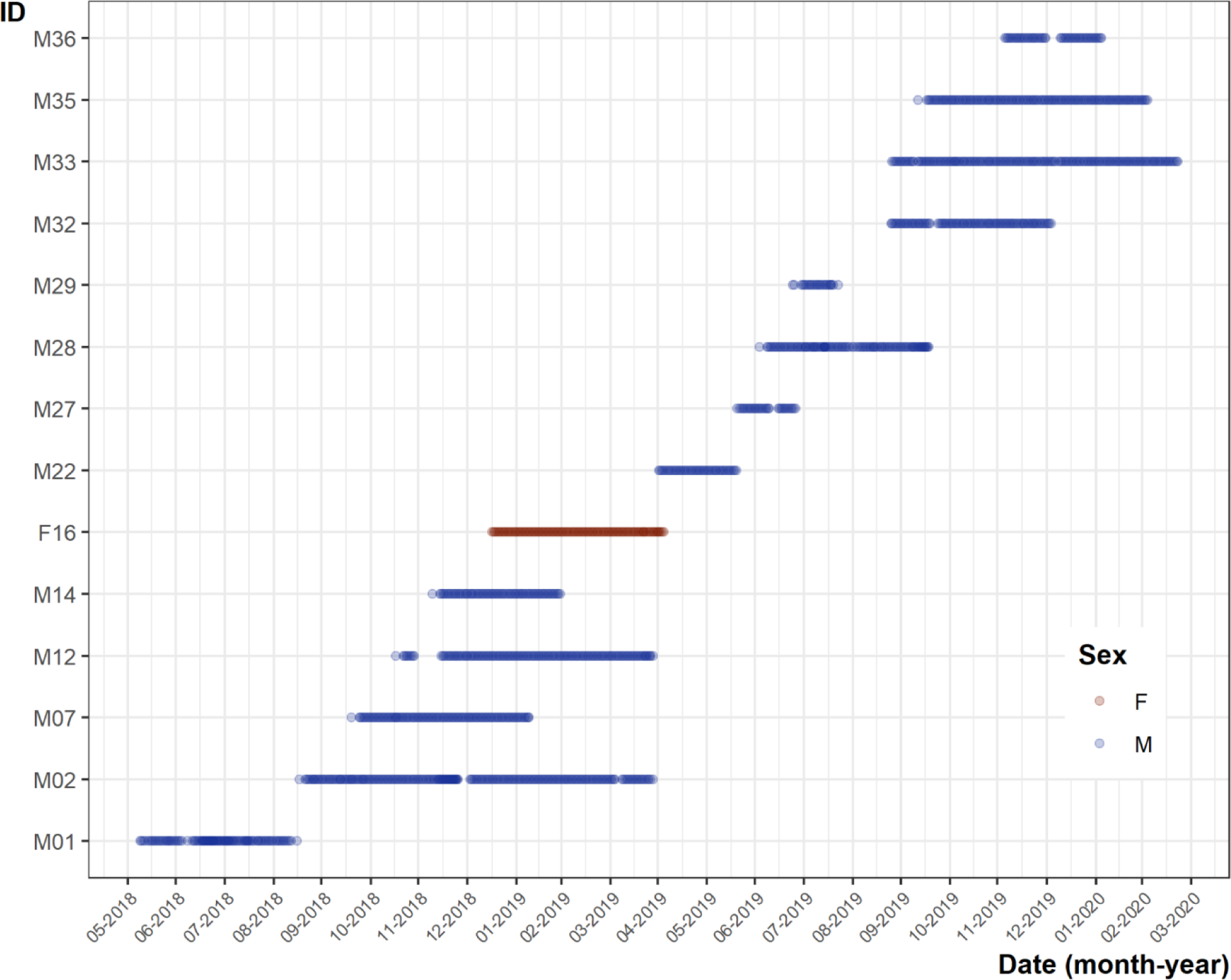
Completed location checks throughout the study period for males (semi-transparent blue) and the female (semi-transparent red), illustrating tracking durations and overlap of simultaneously tracked *B. candidus*.

**Supp. Figure 2.**
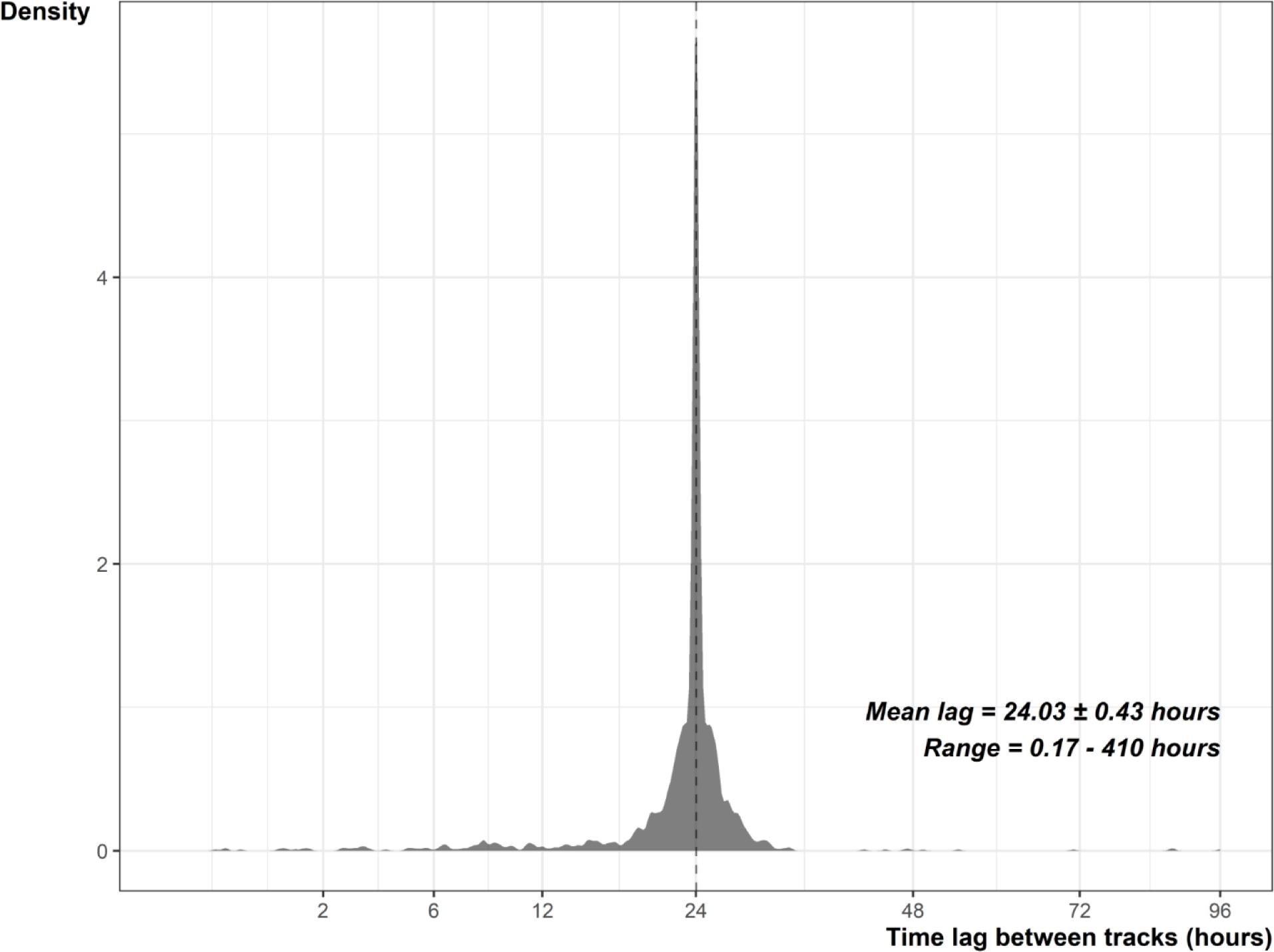
Density plot illustrating time-lags between tracks for all telemetered individuals.

**Supp. Figure 3.**
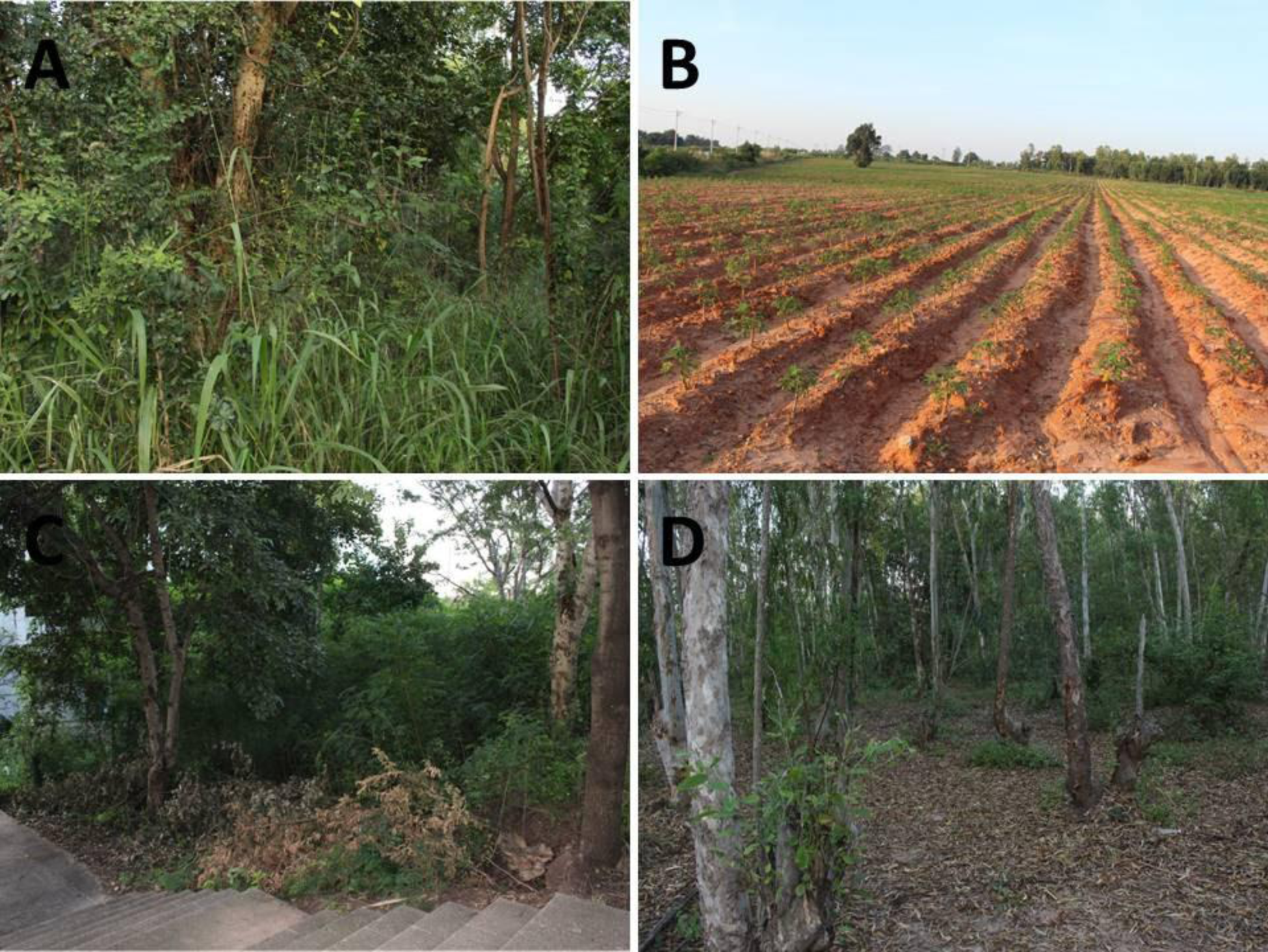
Photos of common land-use types within the study site a) Mixed deciduous forest b) Agriculture (cassava) c) Semi-natural area d) Plantation forest (Eucalyptus)

**Supp. Figure 4.**
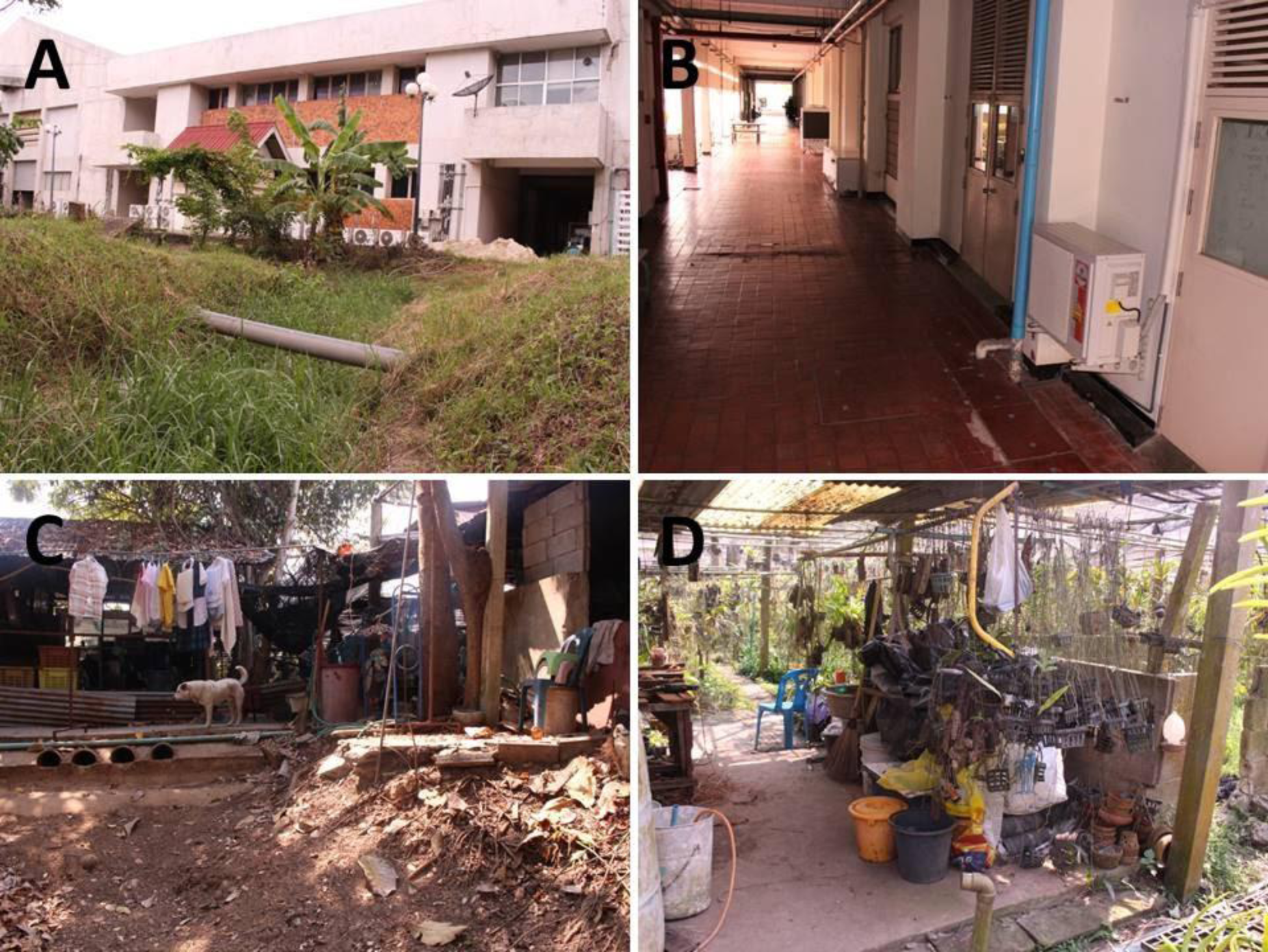
Examples *B. candidus* shelter sites among settlements a) Overgrown concrete drainage ditch outside a laboratory building b) Underneath the foundation or among the pipelines under a laboratory building c) Under a concrete at a residency d) Underneath a concrete floor of a gardening work-station

**Supp. Figure 5.**
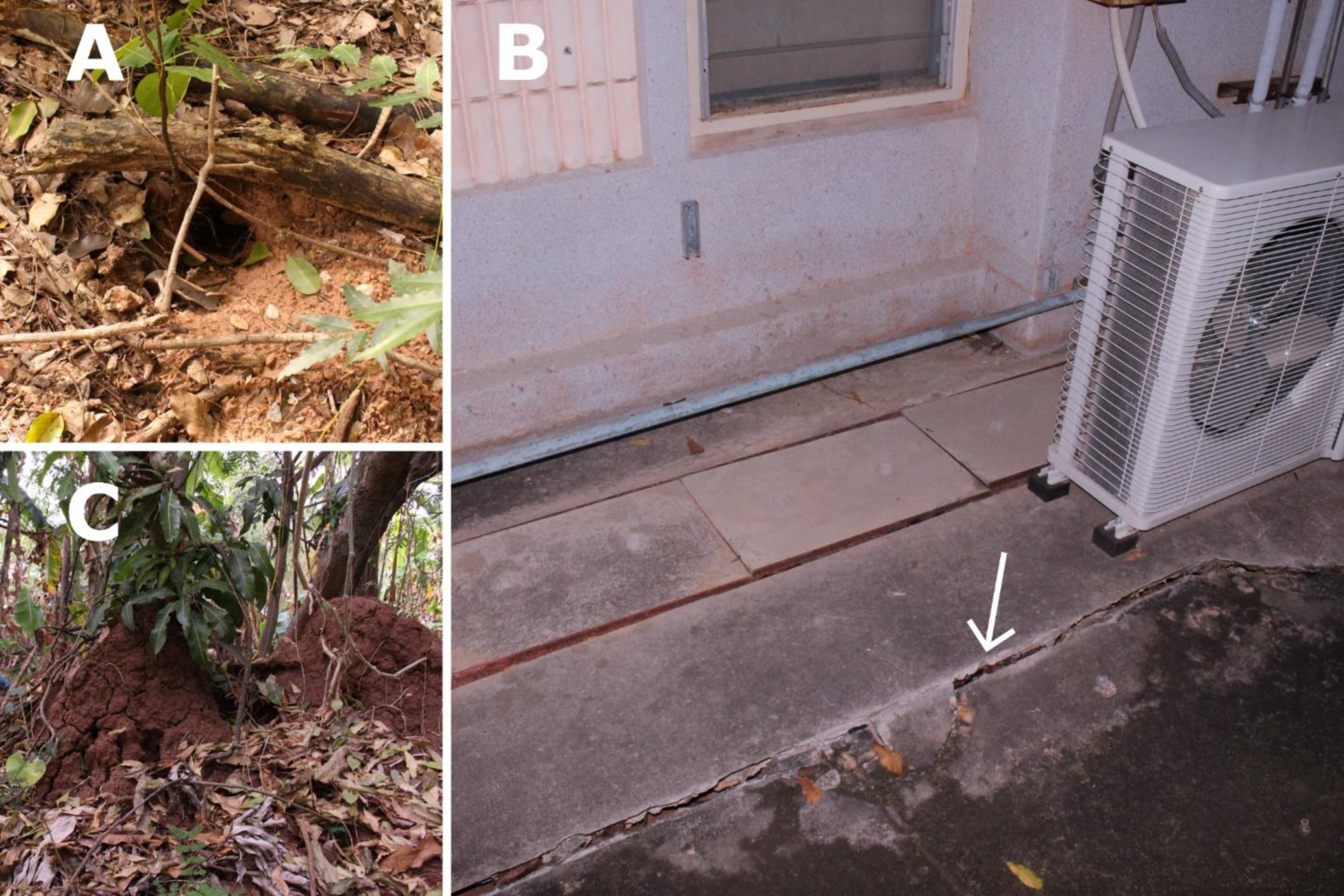
Examples of different shelter types most commonly used by *B. candidus* a) Burrow b) Anthropogenic - under concrete c) Termite mound

**Supp. Figure 6.**
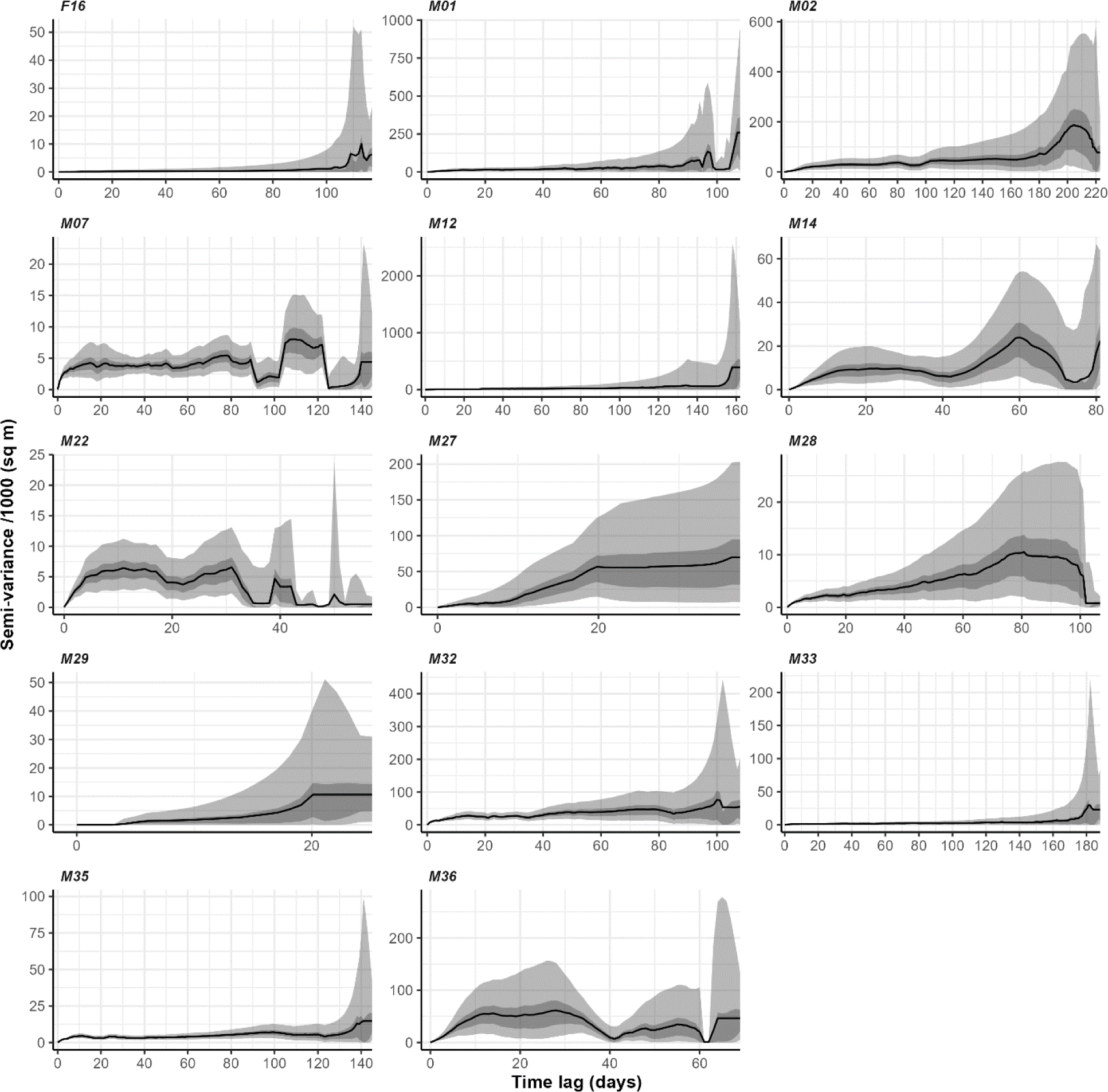
Variogram plots for AKDEs.

**Supp. Figure 7.**
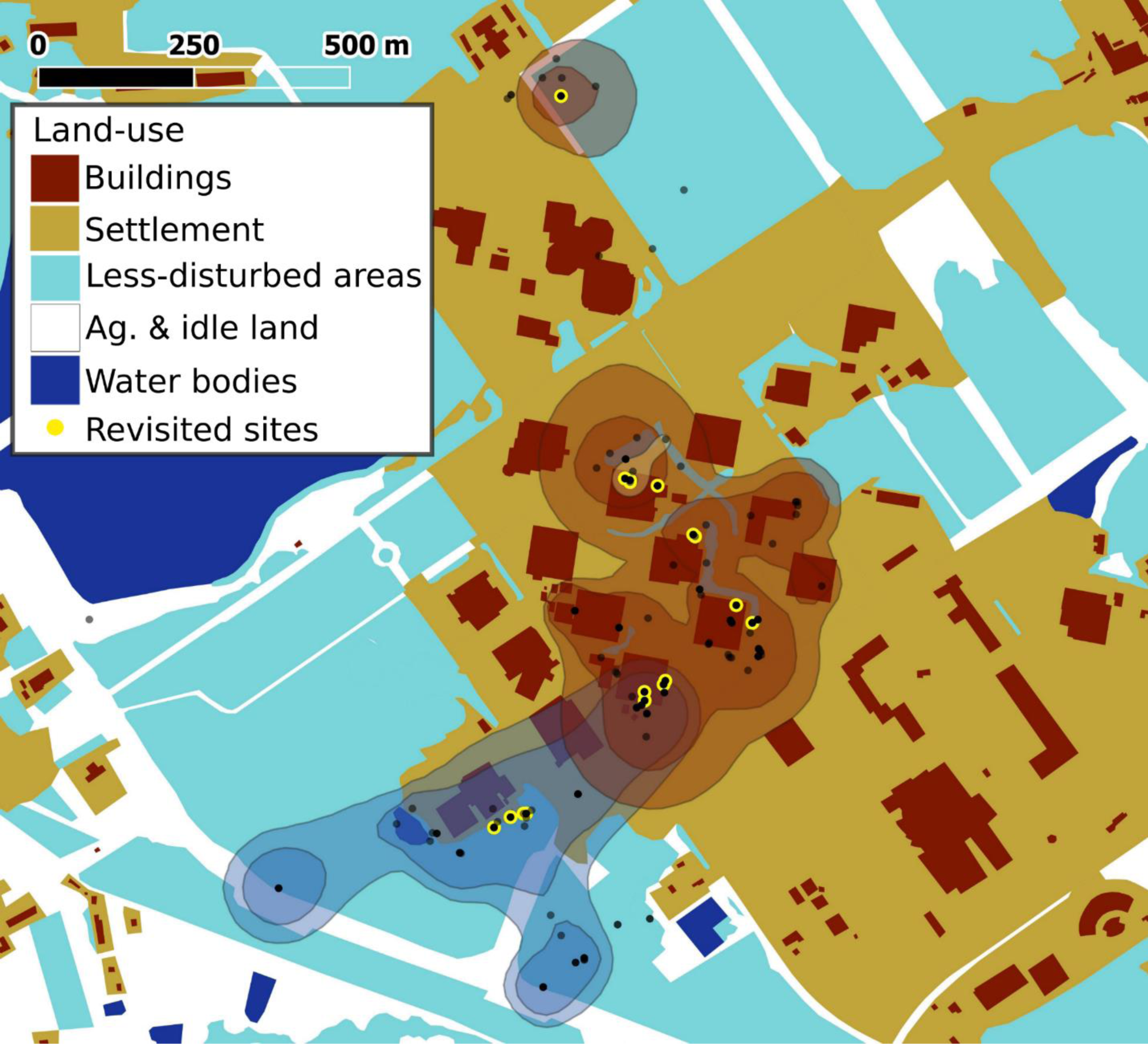
95% and 99% dBBMM occurrence distribution confidence areas for M01 (blue), M02 (red), and F16 (tan) with revisited sites marked in yellow.

**Supp. Figure 8.**
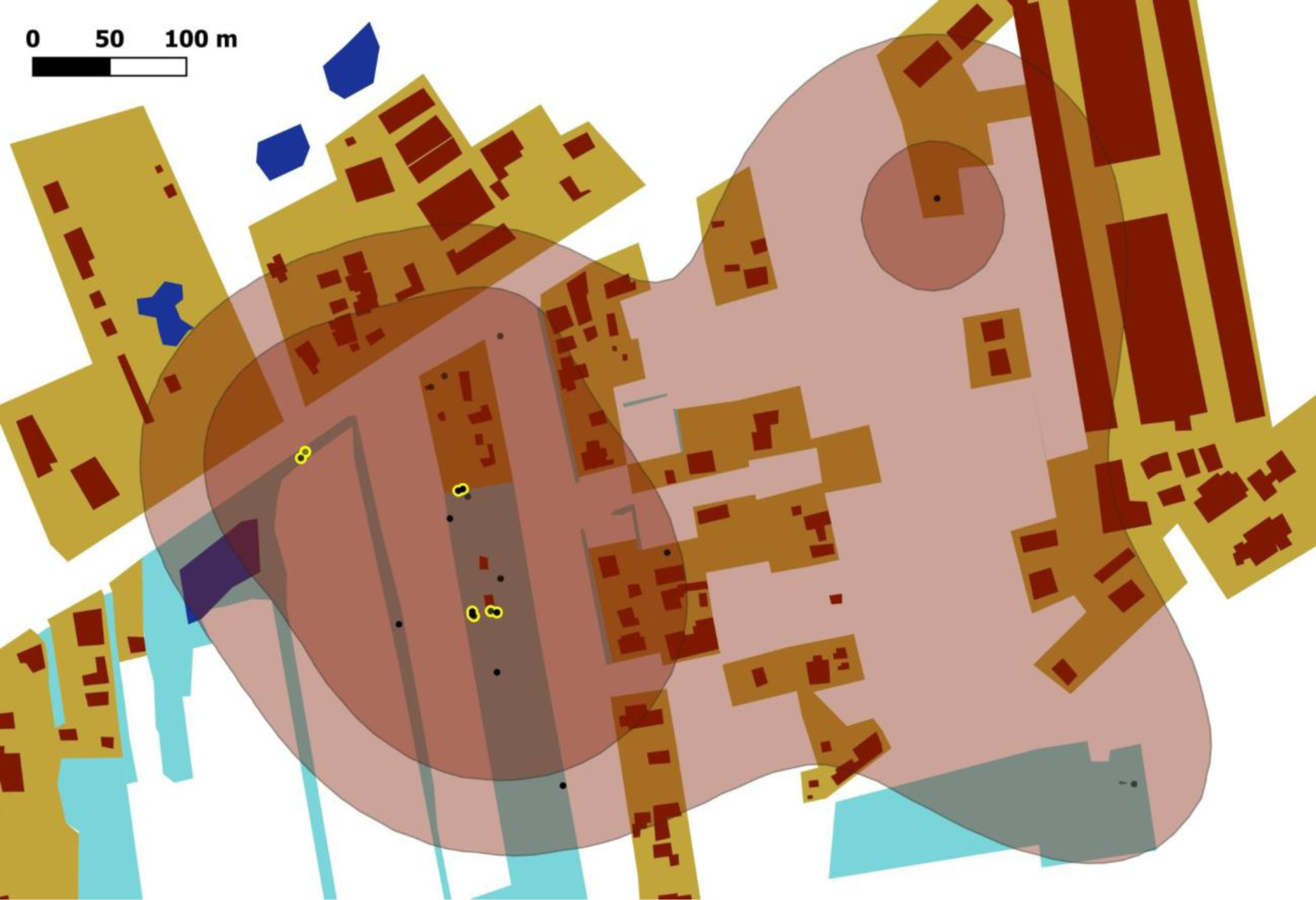
95% and 99% dBBMM occurrence distribution confidence areas for M07 (red) with revisited sites marked in yellow.

**Supp. Figure 9.**
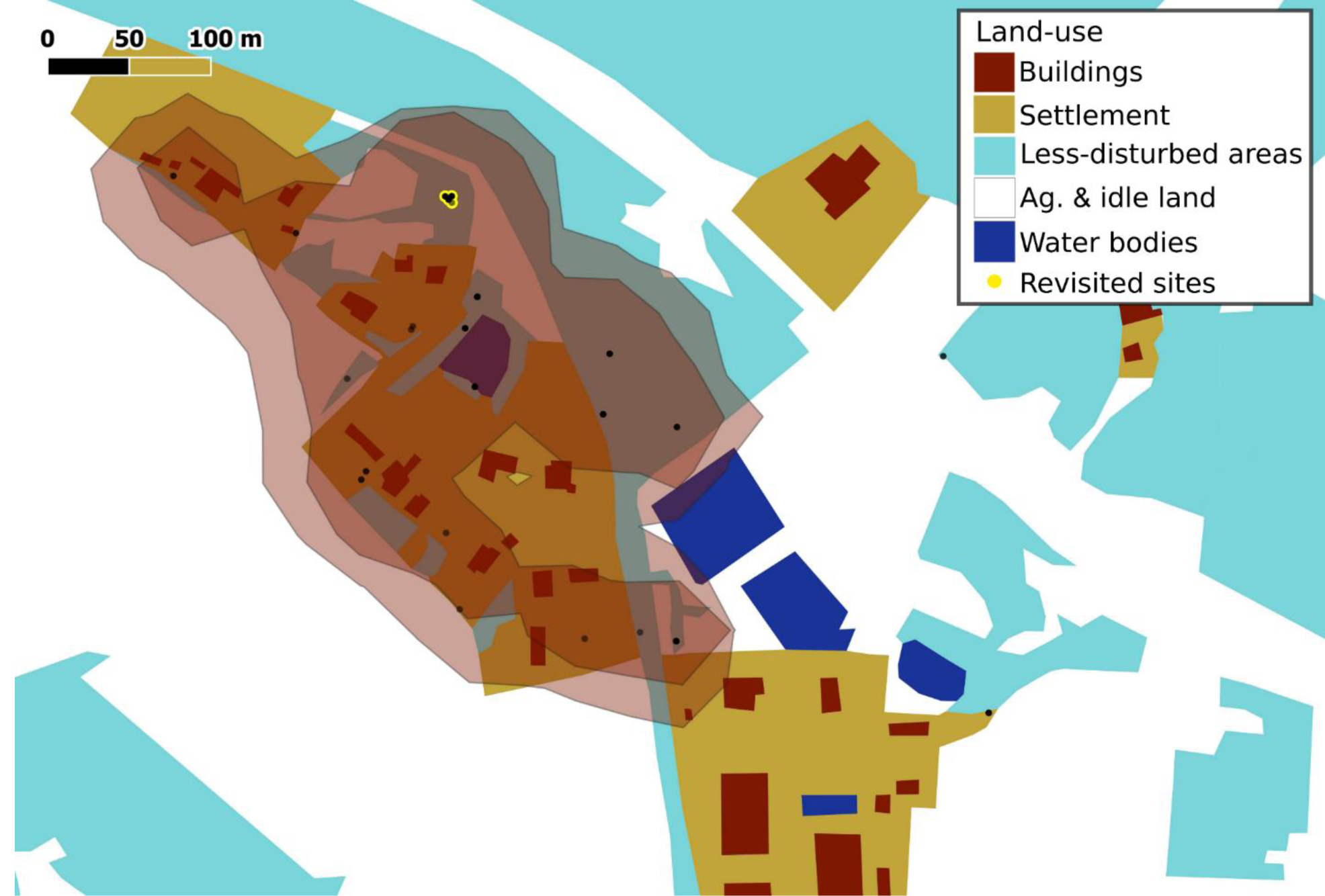
95% and 99% dBBMM occurrence distribution confidence areas for M12 (red) with revisited sites marked in yellow.

**Supp. Figure 10.**
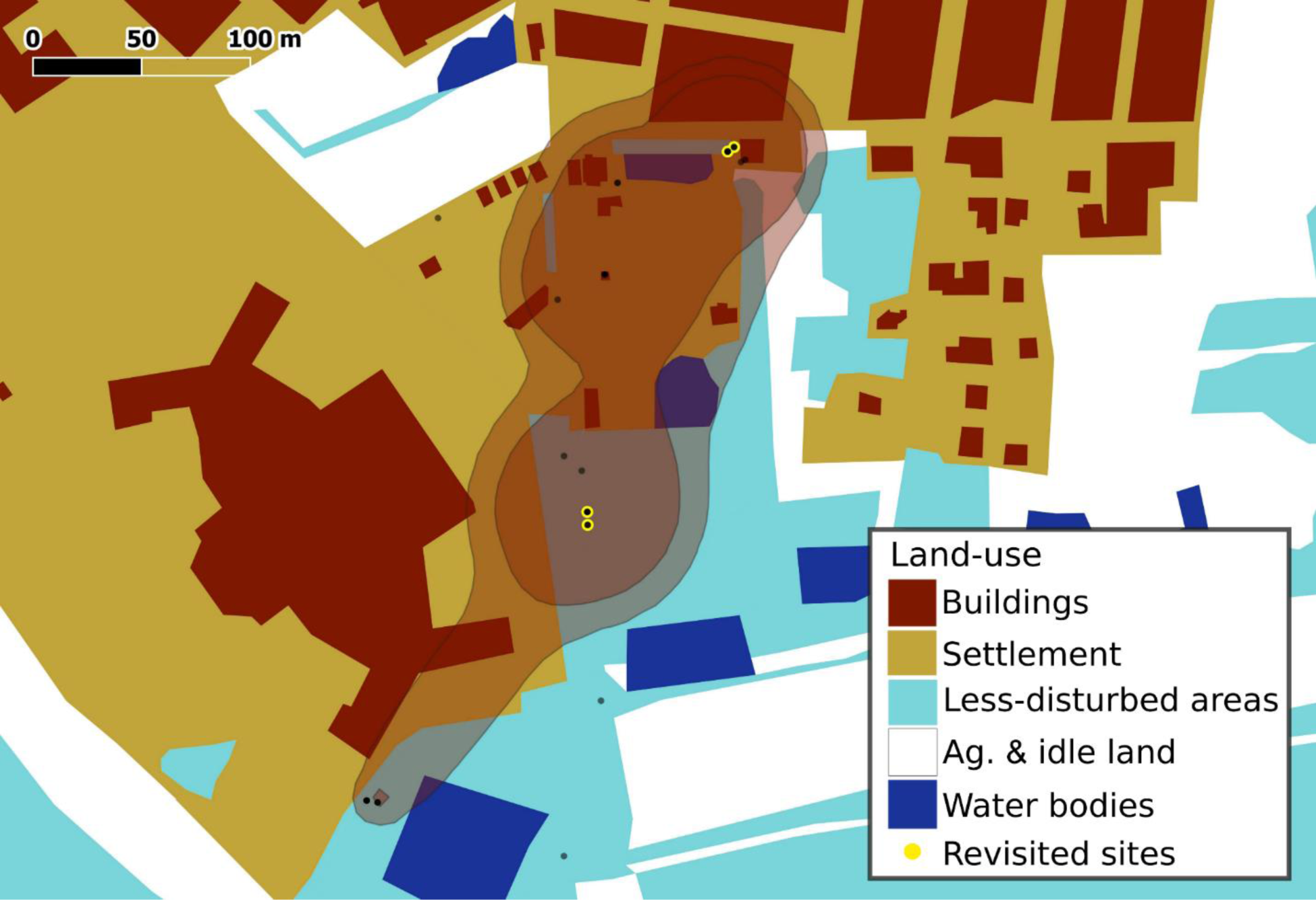
95% and 99% dBBMM occurrence distribution confidence areas for M14 (red) with revisited sites marked in yellow.

**Supp. Figure 11.**
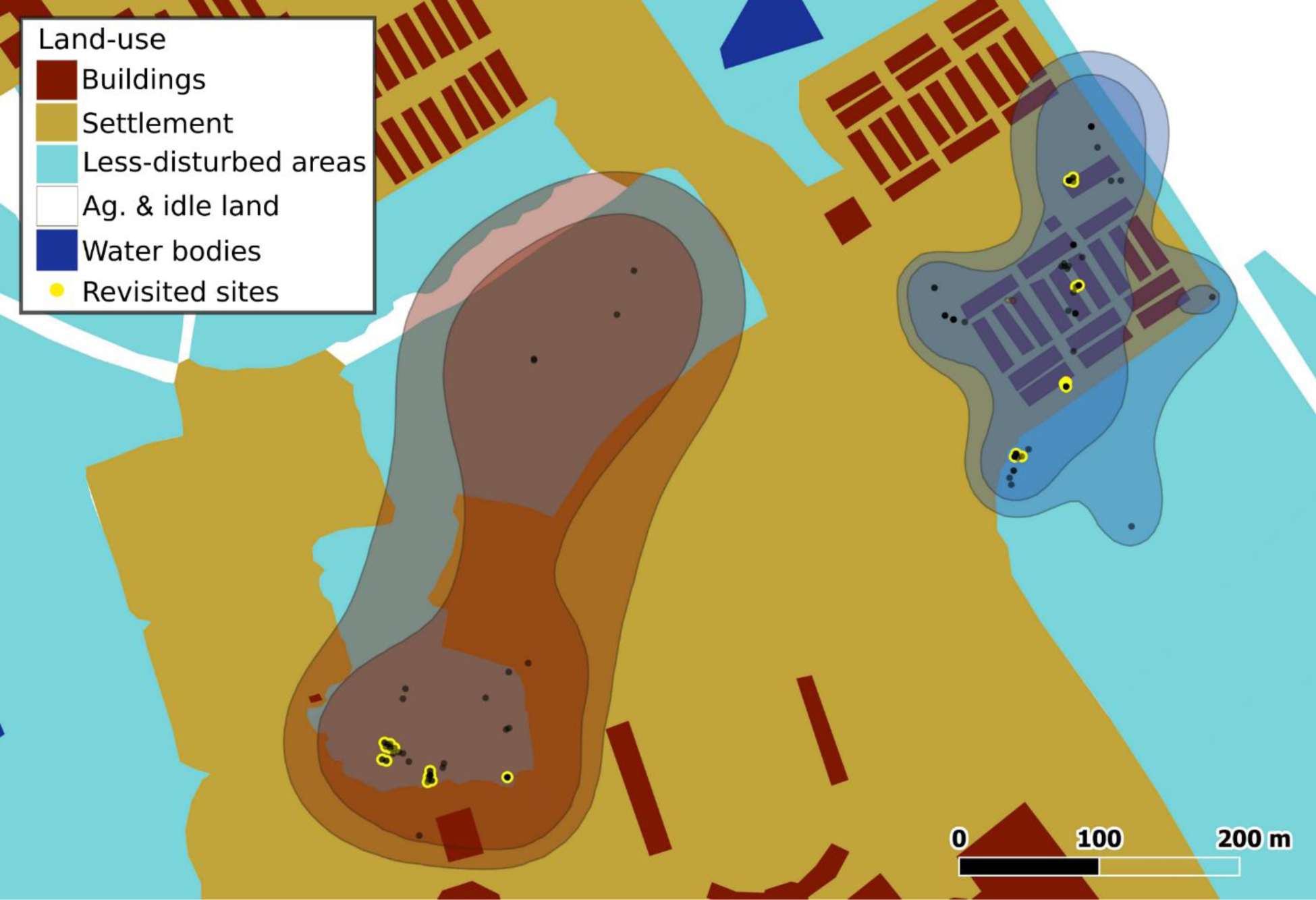
95% and 99% dBBMM occurrence distribution confidence areas for M22 (red) and M28 (blue) with revisited sites marked in yellow.

**Supp. Figure 12.**
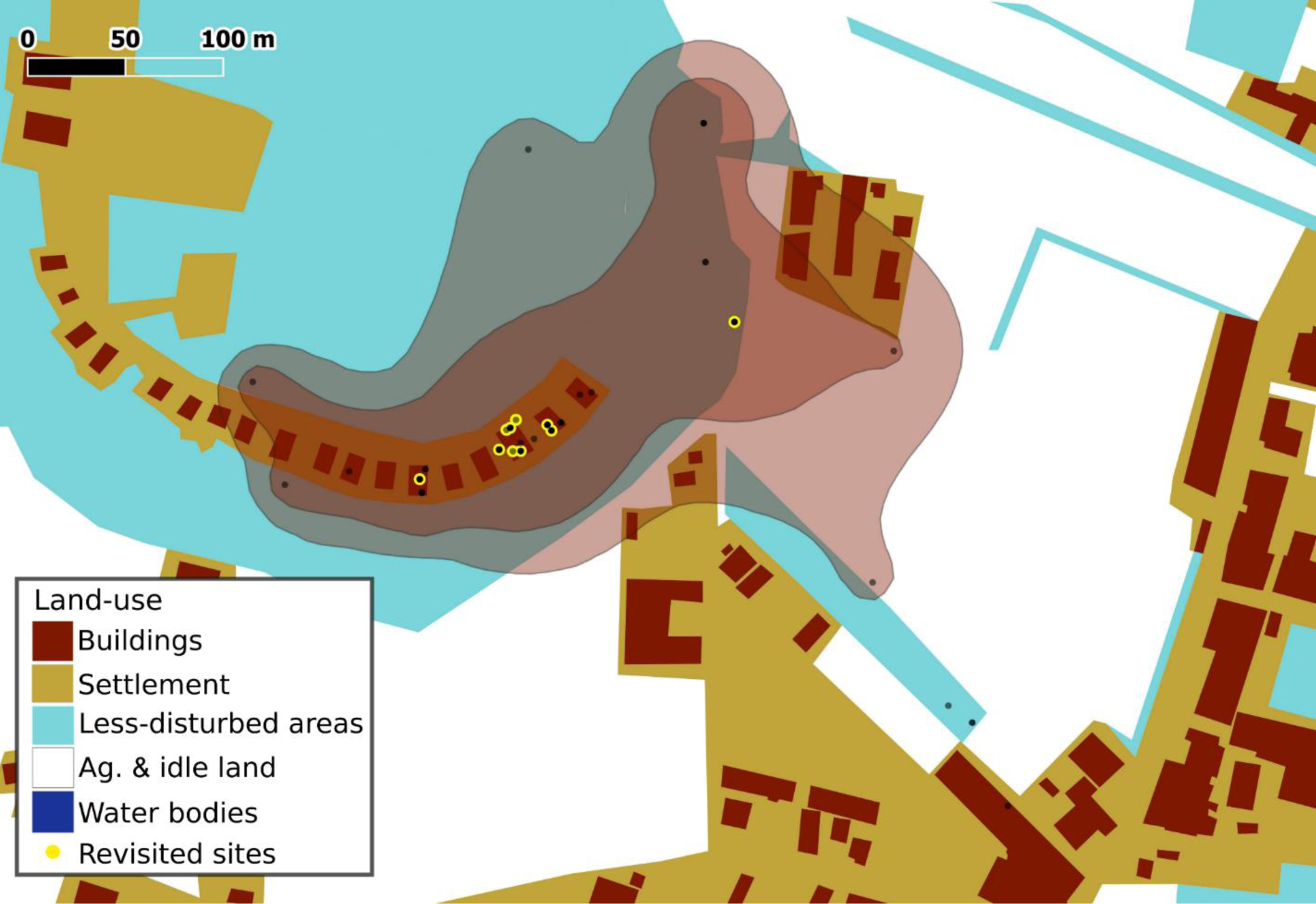
95% and 99% dBBMM occurrence distribution confidence areas for M33 (red) with revisited sites marked in yellow.

**Supp. Figure 13.**
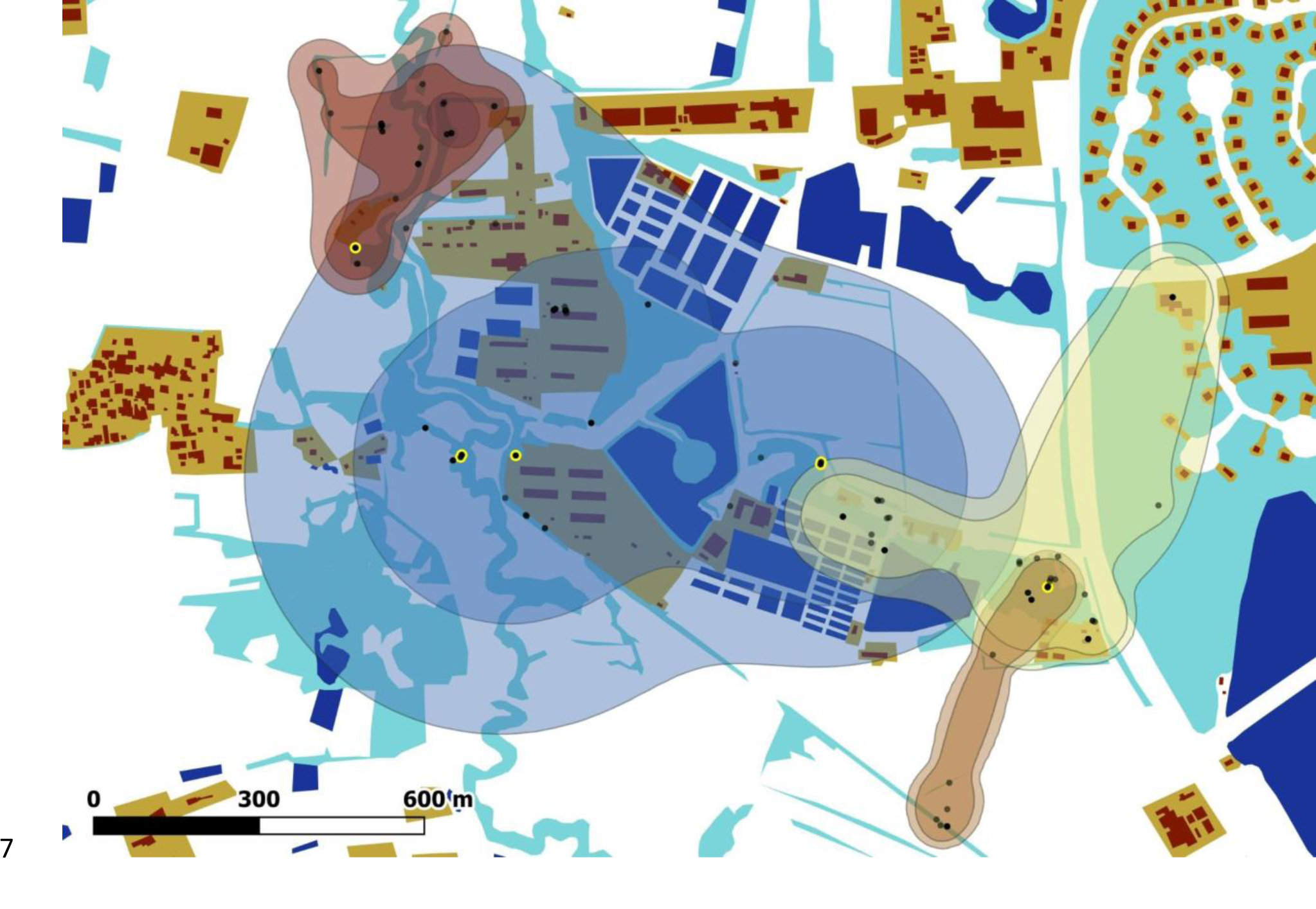
95% and 99% dBBMM occurrence distribution confidence areas for M27 (orange), M32 (blue), M35 (red), and M36 (yellow) with sites revisited marked in yellow.

**Supp. Figure 14.**
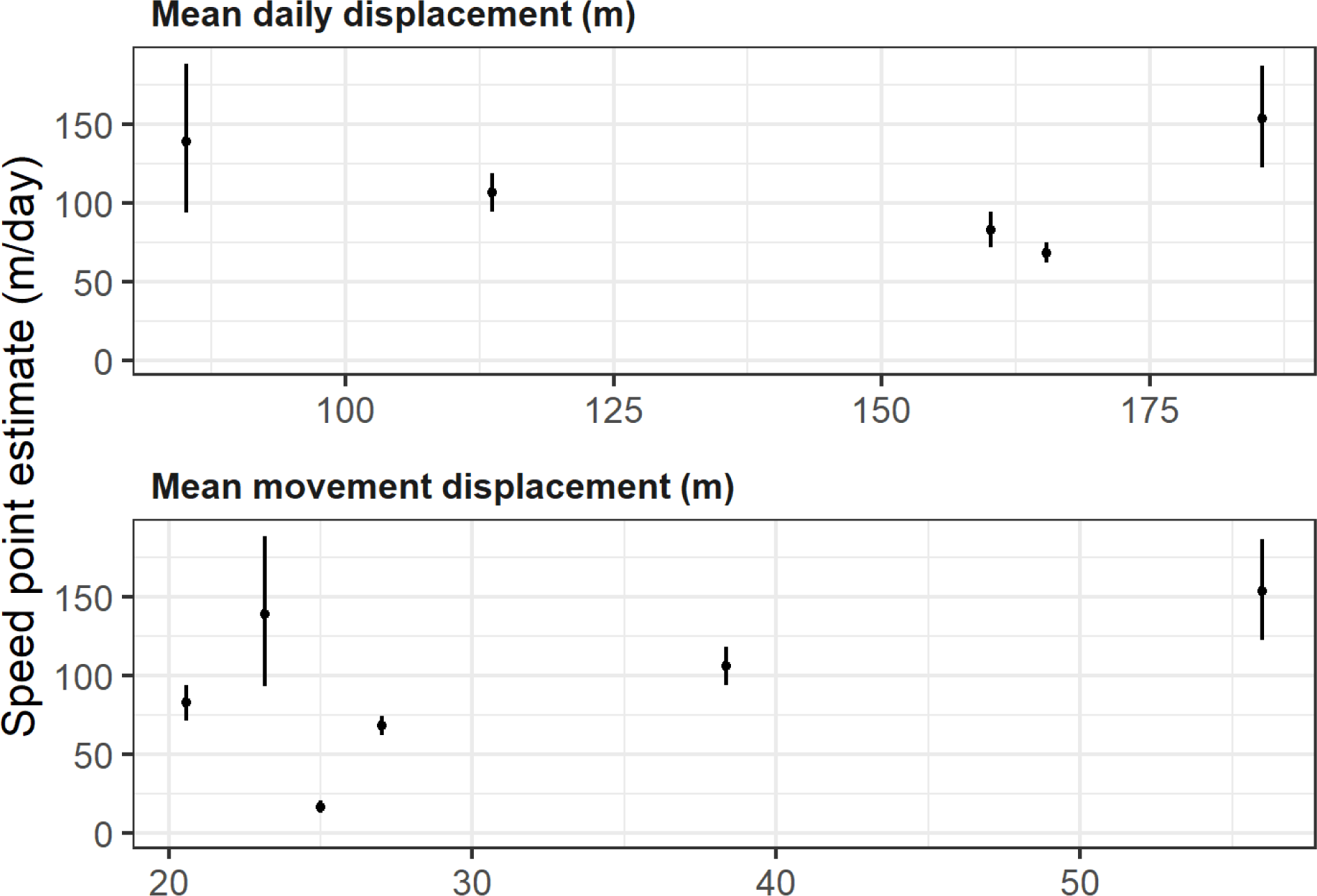
The relationship between mean daily displacement and mean movement displacement and the speed estimates extracted from movement models. Error bars show the 95% CI surrounding the speed estimates. Not all individuals could have speed estimated because of the failure to successfully fit movement models.

**Supp. Figure 15.**
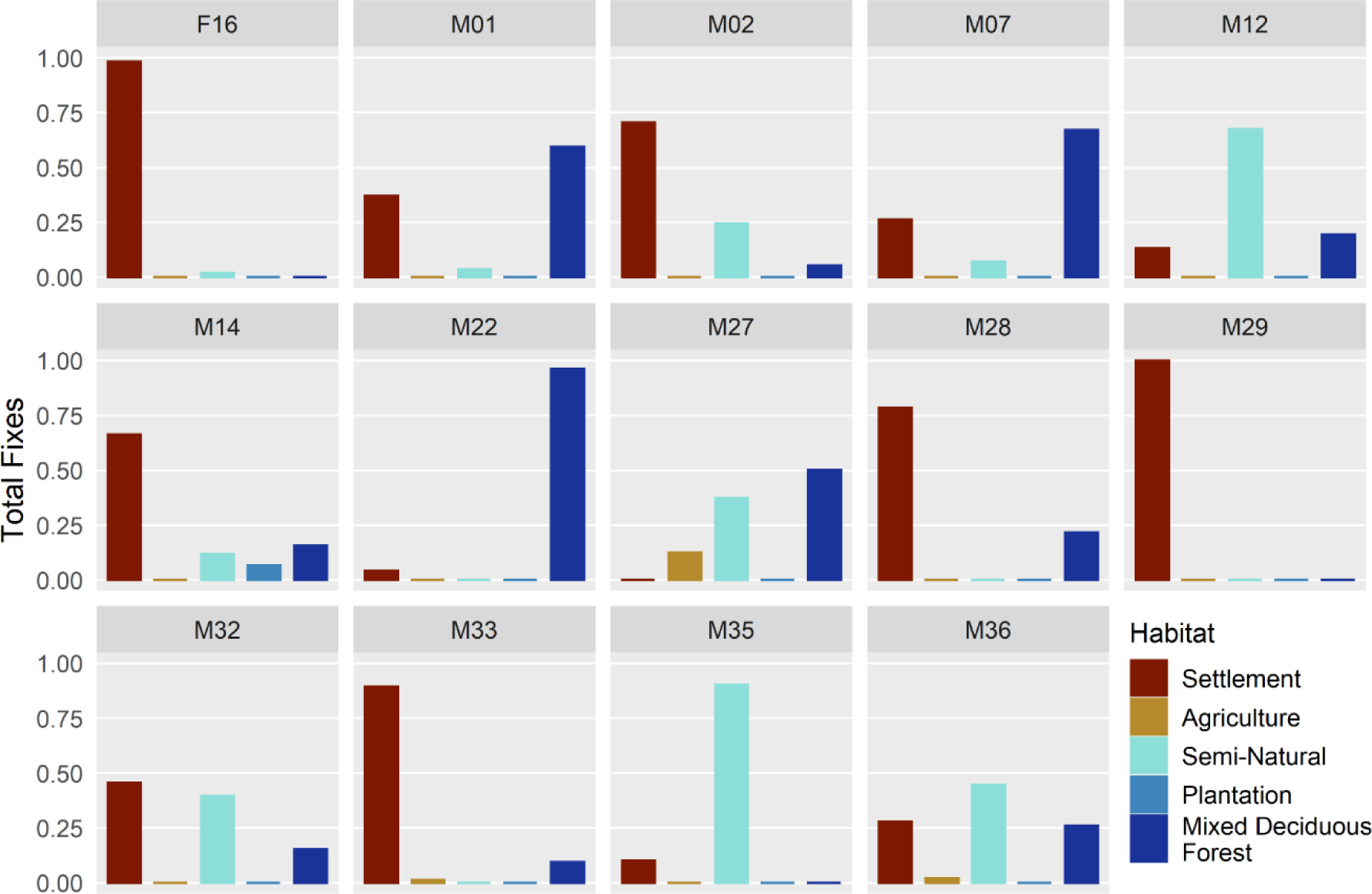
Habitat use proportions for each telemetered individual

**Supp. Figure 16.**
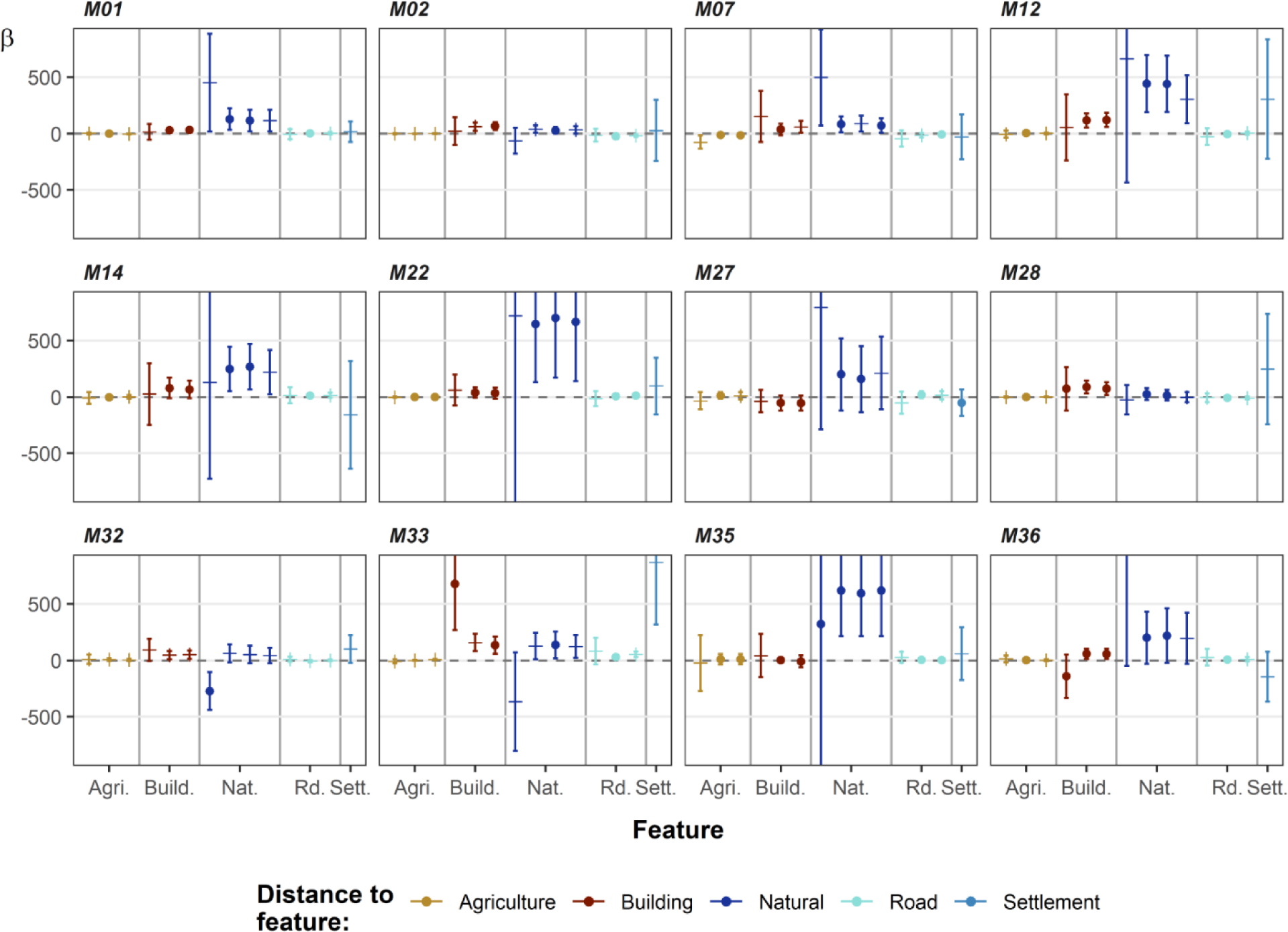
Individual ISSF model results based on distance to habitat features. Positive estimates suggest association with habitat feature, error bars indicate 95% confidence intervals, and circles mark the habitat features that were included in models with AIC scores within <2 D AIC of top performing models.

**Supp. Figure 17.**
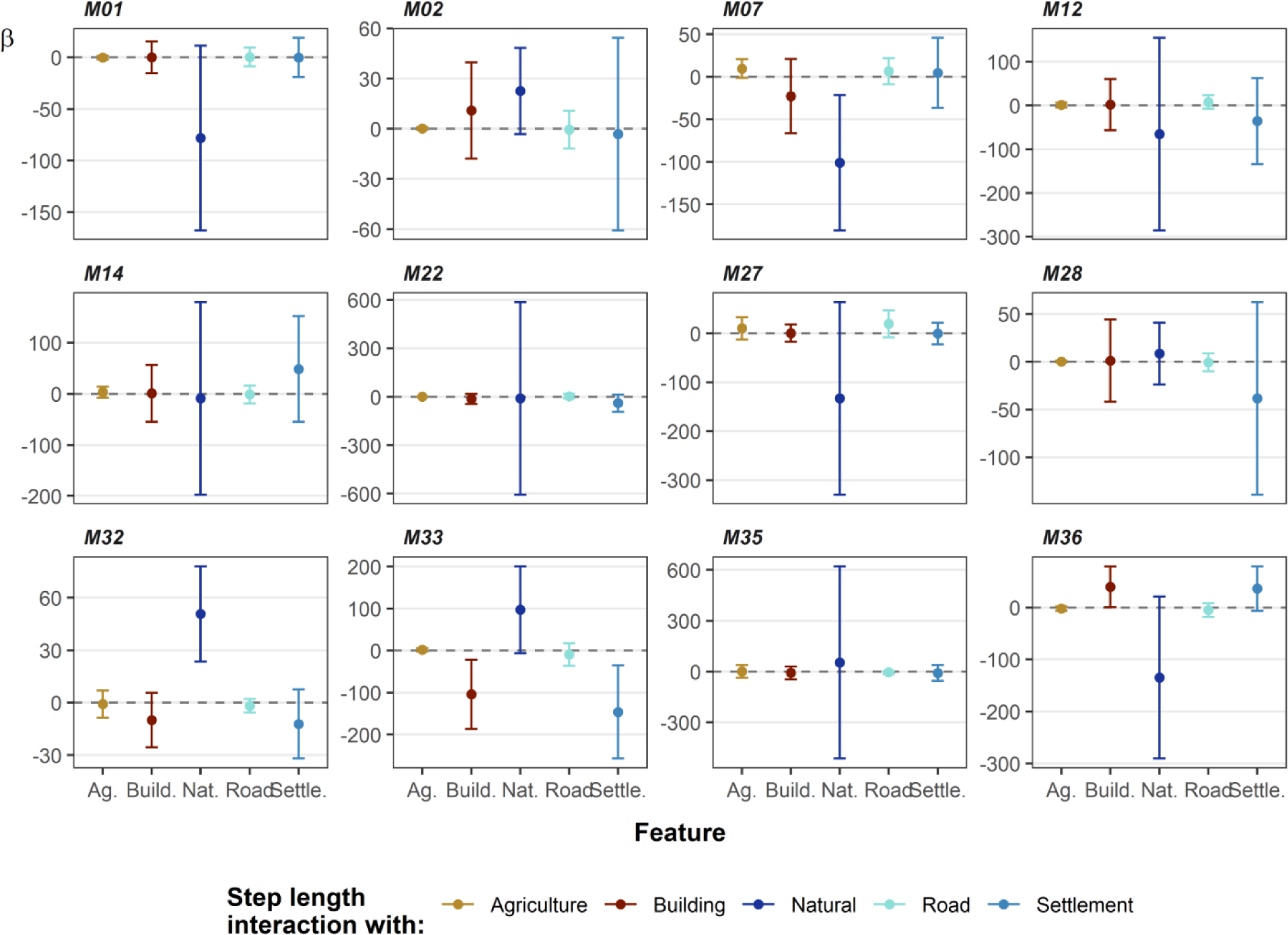
Individual ISSF model results based on the interaction between land-use features and step length. Error bars indicate 95% confidence intervals

**Supp. Table 1.**
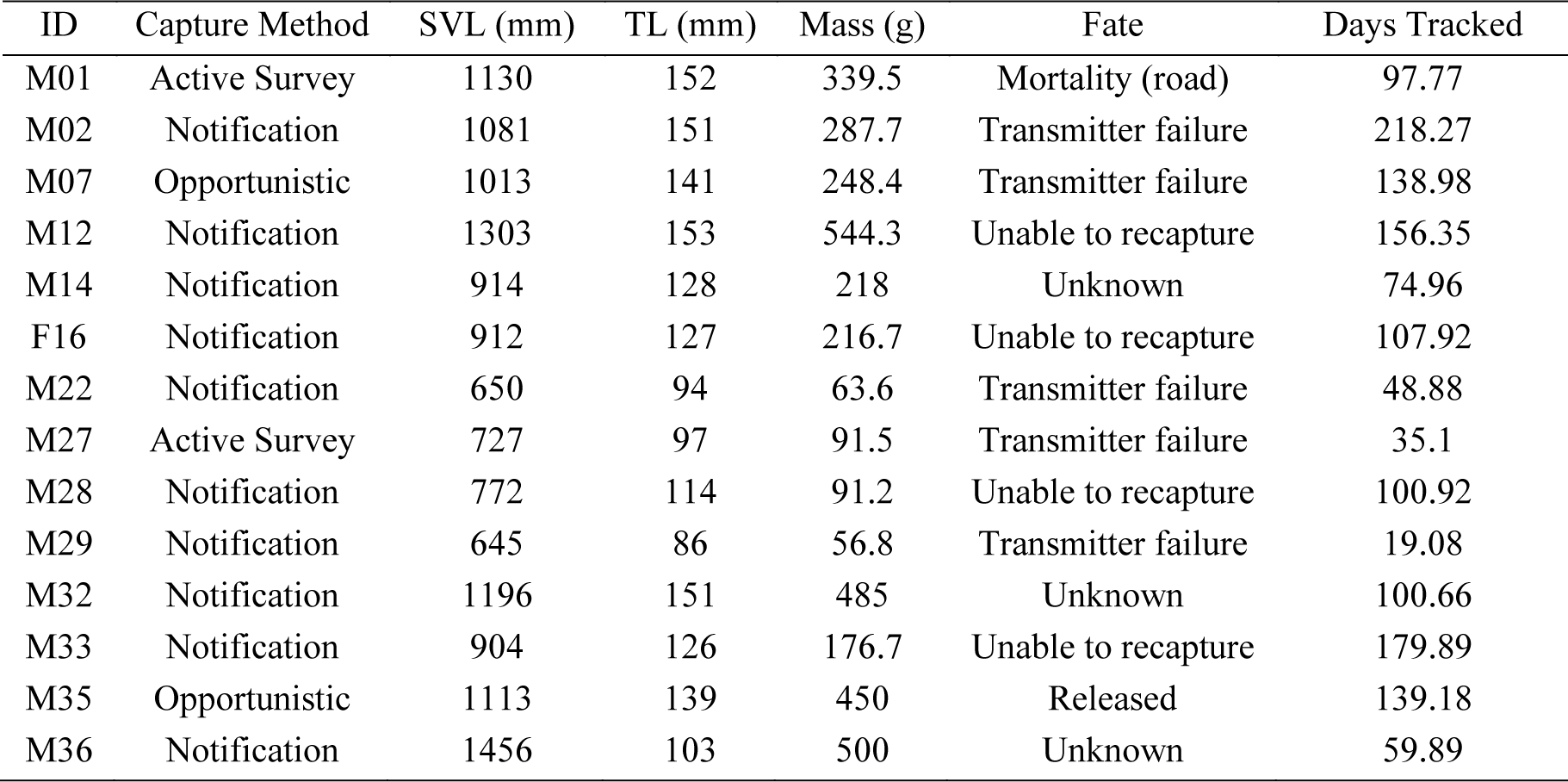
Morphometric data, capture method, days tracked, and fate for each telemetered *B. candidus* individual

| ID | Capture Method | SVL (mm) | TL (mm) | Mass (g) | Fate | Days Tracked |
| --- | --- | --- | --- | --- | --- | --- |
| M01 | Active Survey | 1130 | 152 | 339.5 | Mortality (road) | 97.77 |
| M02 | Notification | 1081 | 151 | 287.7 | Transmitter failure | 218.27 |
| M07 | Opportunistic | 1013 | 141 | 248.4 | Transmitter failure | 138.98 |
| M12 | Notification | 1303 | 153 | 544.3 | Unable to recapture | 156.35 |
| M14 | Notification | 914 | 128 | 218 | Unknown | 74.96 |
| F16 | Notification | 912 | 127 | 216.7 | Unable to recapture | 107.92 |
| M22 | Notification | 650 | 94 | 63.6 | Transmitter failure | 48.88 |
| M27 | Active Survey | 727 | 97 | 91.5 | Transmitter failure | 35.1 |
| M28 | Notification | 772 | 114 | 91.2 | Unable to recapture | 100.92 |
| M29 | Notification | 645 | 86 | 56.8 | Transmitter failure | 19.08 |
| M32 | Notification | 1196 | 151 | 485 | Unknown | 100.66 |
| M33 | Notification | 904 | 126 | 176.7 | Unable to recapture | 179.89 |
| M35 | Opportunistic | 1113 | 139 | 450 | Released | 139.18 |
| M36 | Notification | 1456 | 103 | 500 | Unknown | 59.89 |

**Supp. Table 2.** Table showing capture methods, capture dates, capture locations, release dates, release locations, and translocation distances upon release for every capture and implantation for each telemetered individual.

| ID | Capture date | Capture Easting | Capture Northing | UTM Zone | Capture Method | Release date | Release Easting | Release Northing | UTM Zone | Distance (m) |
| --- | --- | --- | --- | --- | --- | --- | --- | --- | --- | --- |
| M01 | 05-30-2018<br>22:59 | 179060 | 1646306 | 48 | Active Survey | 06-08-2018<br>19:44 | 179060 | 1646306 | 48 | 0 |
| M01 | 09-11-2018<br>19:30 | 178813 | 1646489 | 48 | Tracked to capture | 09-12-2018<br>21:00 | 178813 | 1646489 | 48 | 0 |
| M02 | 09-17-2018<br>00:00 | 179007 | 1647371 | 48 | Notification | 09-20-2018<br>23:00 | 179093 | 1647380 | 48 | 86.47 |
| M07 | 10-19-2018<br>21:30 | 182020 | 1648494 | 48 | Opportunistic capture | 10-24-2018<br>23:30 | 182020 | 1648494 | 48 | 0 |
| M12 | 11-16-2018<br>19:30 | 181180 | 1644862 | 48 | Notification | 11-21-2018<br>19:20 | 181286 | 1644926 | 48 | 123.82 |
| M14 | 12-09-2018<br>20:13 | 180581 | 1646722 | 48 | Notification | 12-14-2018<br>20:30 | 180636 | 1646611 | 48 | 123.88 |
| M02 | 12-25-2018<br>20:30 | 179062 | 1646598 | 48 | Tracked to capture | 01-02-2019<br>19:30 | 179092 | 1646681 | 48 | 88.26 |
| F16 | 01-08-2019<br>07:20 | 179113 | 1646872 | 48 | Notification | 01-14-2019<br>23:00 | 179166 | 1646874 | 48 | 53.04 |
| M07 | 02-08-2019<br>12:30 | 181904 | 1648459 | 48 | Tracked to capture | 02-21-2019<br>23:30 | 181892 | 1648457 | 48 | 12.17 |
| M02 | 04-03-2019<br>20:08 | 179228 | 1646700 | 48 | Tracked to capture | 04-07-2019<br>22:30 | 179228 | 1646700 | 48 | 0 |
| M22 | 04-23-2019<br>17:45 | 178776 | 1648029 | 48 | Notification | 04-30-2019<br>22:00 | 178783 | 1648067 | 48 | 38.64 |
| M27 | 06-17-2019<br>20:25 | 177709 | 1647409 | 48 | Active Survey | 06-20-2019<br>21:00 | 177709 | 1647409 | 48 | 0 |
| M28 | 07-03-2019<br>22:10 | 179260 | 1648429 | 48 | Notification | 07-08-2019<br>23:40 | 179210 | 1648287 | 48 | 150.55 |
| M27 | 07-09-2019<br>21:15 | 177496 | 1646956 | 48 | Tracked to capture | 07-15-2019<br>22:30 | 177476 | 1646938 | 48 | 26.91 |
| M29 | 07-25-2019<br>23:50 | 180146 | 1646557 | 48 | Notification | 07-30-2019<br>00:15 | 180338 | 1646437 | 48 | 226.42 |
| M32 | 09-17-2019<br>23:50 | 822770 | 1647515 | 47 | Notification | 09-24-2019<br>20:40 | 822817 | 1647592 | 47 | 90.21 |
| M33 | 09-18-2019<br>00:12 | 178254 | 1649809 | 48 | Notification | 09-24-2019<br>21:20 | 178225 | 1649861 | 48 | 59.54 |
| M33 | 10-08-2019<br>20:25 | 178202 | 1650043 | 48 | Tracked to capture | 10-11-2019<br>23:50 | 178121 | 1650060 | 48 | 82.76 |
| M35 | 10-11-2019<br>19:30 | 822301 | 1648030 | 47 | Opportunistic capture | 10-15-2019<br>23:59 | 822344 | 1648028 | 47 | 43.05 |
| M33 | 11-03-2019<br>21:39 | 178188 | 1649925 | 48 | Tracked to capture | 11-04-2019<br>19:40 | 178121 | 1650060 | 48 | 150.71 |
| M36 | 11-27-2019<br>20:42 | 177371 | 1647443 | 48 | Notification | 12-05-2019<br>18:40 | 177372 | 1647458 | 48 | 15.03 |

**Supp. Table 3.** Movement and space use summary data for each telemetered *B. candidus*. Dynamic Brownian Bridge Movement model 90%, 95%, and 99% confidence areas (hectares), mean motion variance (σ²m), mean movement distance (MMD, in meters), mean daily displacement (MDD, in meters), and relocation probabilities (i.e., proportion of daily fixes moved) shown with overall averages for males (n = 13) (note M29 excluded from dBBMM occurrence distribution and motion variance means, n = 12).

| ID | 90%<br>(ha) | 95%<br>(ha) | 99%<br>(ha) | $\sigma^2m$ | MMD (m) | MDD (m) | Move<br>probability |
| --- | --- | --- | --- | --- | --- | --- | --- |
| M01 | 6.73 | 10.57 | 18.8 | $1.23 \pm 0.17$ | $113.7 \pm 26.51$ | $38.35 \pm 10.53$ | 0.34 |
| M02 | 7.7 | 12.91 | 25.68 | $1.2 \pm 0.19$ | $85.18 \pm 12.2$ | $23.17 \pm 4.20$ | 0.27 |
| M07 | 2.16 | 5.58 | 23.39 | $2.37 \pm 0.53$ | $132.87 \pm 25.14$ | $24.11 \pm 6.38$ | 0.18 |
| M12 | 3.11 | 5.04 | 8.4 | $0.66 \pm 0.08$ | $165.39 \pm 46.02$ | $27.01 \pm 9.0$ | 0.16 |
| M14 | 1.06 | 1.52 | 2.52 | $0.55 \pm 0.06$ | $93.53 \pm 16.76$ | $21.03 \pm 5.8$ | 0.22 |
| F16 | 0.03 | 0.05 | 0.42 | $0.16 \pm 0.04$ | $47.15 \pm 11.75$ | $5.68 \pm 2.0$ | 0.12 |
| M22 | 3.96 | 6.05 | 10.07 | $2.33 \pm 0.34$ | $69.85 \pm 16.03$ | $37.0 \pm 9.6$ | 0.52 |
| M27 | 2.79 | 3.63 | 5.33 | $0.6 \pm 0.2$ | $80.86 \pm 33.93$ | $19.62 \pm 10.11$ | 0.24 |
| M28 | 1.89 | 2.77 | 4.81 | $0.58 \pm 0.09$ | $66.45 \pm 10.68$ | $19.22 \pm 4.26$ | 0.29 |
| M29 | 0.03 | 0.04 | 0.06 | na | $91.67 \pm 79.67$ | $25.0 \pm 22.65$ | 0.27 |
| M32 | 29.8 | 56.2 | 119.55 | $9.63 \pm 2.34$ | $259.65 \pm 57.67$ | $60.99 \pm 17.37$ | 0.23 |
| M33 | 1.61 | 3.11 | 6.46 | $0.7 \pm 0.11$ | $82.77 \pm 10.44$ | $14.54 \pm 2.98$ | 0.18 |
| M35 | 3.39 | 6.79 | 13.09 | $1.25 \pm 0.21$ | $160.17 \pm 19.66$ | $20.59 \pm 5.17$ | 0.13 |
| M36 | 15.71 | 20.47 | 29.85 | $4.38 \pm 0.52$ | $185.44 \pm 46.23$ | $55.98 \pm 18.04$ | 0.3 |
| Avg<br>(males) | $6.66 \pm 2.41$ | $11.22 \pm 4.37$ | $22.33 \pm 9.21$ | $1.70 \pm 0.18$ | $117.78 \pm 8.23$ | $27.48 \pm 2.36$ | 0.23 |

**Supp. Table 4.** Results from AKDE analysis, including effective sample size (DOF_area), the type of top fitted model (movMod), the 95% contour AKDE area estimates (ha) with 95% CI, the speed estimate with 95% CI is calculable for the top movement model (m/day).

| ID | DOF<br>area | Move model | aKDE<br>low | aKDE est | aKDE<br>high | Speed<br>low | Speed est | Speed<br>high |
| --- | --- | --- | --- | --- | --- | --- | --- | --- |
| F16 | 3.87 | OU anisotropic | 0.62 | 2.33 | 5.17 | 0 | Inf | Inf |
| M01 | 3.06 | OUF anisotropic | 17.29 | 82.07 | 196.32 | 94.3 | 106.36 | 118.77 |
| M02 | 5.49 | OUF anisotropic | 22.85 | 65.94 | 131.45 | 93.58 | 138.94 | 188.47 |
| M07 | 55.22 | OU isotropic | 4.03 | 5.34 | 6.84 | 0 | Inf | Inf |
| M12 | 1.49 | OUF anisotropic | 0.35 | 4.99 | 15.58 | 62.19 | 68.36 | 74.68 |
| M14 | 2.19 | OU anisotropic | 2.52 | 18.19 | 48.99 | 0 | Inf | Inf |
| M22 | 19.98 | OU anisotropic | 2.7 | 4.42 | 6.56 | 0 | Inf | Inf |
| M27 | 1.87 | OU anisotropic | 1.51 | 13.93 | 39.83 | 0 | Inf | Inf |
| M28 | 135.75 | IID anisotropic | 2.75 | 3.28 | 3.86 | 0 | Inf | Inf |
| M29 | 1.9 | Ouf anisotropic | 0.54 | 4.83 | 13.73 | 12.85 | 16.76 | 20.92 |
| M32 | 14.12 | OU anisotropic | 35.7 | 65.1 | 103.19 | 0 | Inf | Inf |
| M33 | 15.56 | OU isotropic | 3.9 | 6.89 | 10.7 | 0 | Inf | Inf |
| M35 | 39.38 | OUF anisotropic | 5.46 | 7.67 | 10.24 | 71.68 | 82.81 | 94.32 |
| M36 | 5.65 | OUF anisotropic | 31.16 | 88.23 | 174.49 | 122.63 | 153.94 | 186.92 |

**Supp. Table 5.** Total number of fixes an individual was located in each land-use type. MDF = mixed deciduous forest.

| Snake ID | Settlement | Semi-Nat | Agriculture | MDF | Plantation |
| --- | --- | --- | --- | --- | --- |
| M01 | 43 | 4 | 0 | 69 | 0 |
| M02 | 162 | 56 | 0 | 12 | 0 |
| M07 | 34 | 9 | 0 | 87 | 0 |
| M12 | 19 | 97 | 0 | 28 | 0 |
| M14 | 51 | 9 | 0 | 12 | 5 |
| F16 | 110 | 2 | 0 | 0 | 0 |
| M22 | 2 | 0 | 0 | 49 | 0 |
| M27 | 0 | 12 | 4 | 16 | 0 |
| M28 | 91 | 0 | 0 | 25 | 0 |
| M29 | 24 | 0 | 0 | 0 | 0 |
| M32 | 45 | 39 | 0 | 15 | 0 |
| M33 | 160 | 0 | 2 | 17 | 0 |
| M35 | 14 | 127 | 0 | 0 | 0 |
| M36 | 15 | 24 | 1 | 14 | 0 |
| *Total* | *770 (51.2%)* | *379 (25.2%)* | *7 (0.5%)* | *344 (22.8%)* | *5 (0.3%)* |

**Supp. Table 6.** Total number of fixes an individual was located within each shelter type

| ID | Burrow | Termite Md | Anthro | Unk |
| --- | --- | --- | --- | --- |
| M01 | 27 | 20 | 40 | 5 |
| M02 | 72 | 12 | 135 | 2 |
| M07 | 116 | 7 | 2 | 3 |
| M12 | 67 | 62 | 8 | 6 |
| M14 | 41 | 1 | 9 | 25 |
| F16 | 0 | 1 | 108 | 1 |
| M22 | 44 | 6 | 0 | 0 |
| M27 | 27 | 0 | 0 | 2 |
| M28 | 70 | 4 | 33 | 0 |
| M29 | 0 | 0 | 20 | 1 |
| M32 | 42 | 1 | 44 | 10 |
| M33 | 23 | 5 | 148 | 0 |
| M35 | 36 | 95 | 0 | 9 |
| M36 | 17 | 3 | 12 | 21 |
| *Total* | *582*<br>*(40.3%)* | *217*<br>*(15%)* | *559*<br>*(38.7%)* | *85*<br>*(5.9%)* |

**Supp. Table 7.** Model formulas and AIC scores for individual ISSF models. * with emboldened text indicates AIC scores within < 2 Δ AIC of the model that best supports the observed movements for an individual

| model<br>formula | M01 | M02 | M07 | M12 | M14 | M22 | M27 | M28 | M32 | M33 | M35 | M36 |
| --- | --- | --- | --- | --- | --- | --- | --- | --- | --- | --- | --- | --- |
| model1<br>(null) | 345.11 | 620.56 | 249.4 | 239.03 | 175.24 | 300.2 | 75.62 | 358 | 259.89 | 310.62 | 185.6 | 163.86 |
| Settlement | 346.5 | 626.17 | 254.49 | 225.74 | 180.15 | 298.2 | 75.7 | 357.73 | 259.84 | 290 | 190.98 | 160.5 |
| Road | 350.5 | 615.23 | 251.22 | 239.16 | 180.62 | 305.21 | 77.42 | 360.64 | 260.81 | 300.81 | 190.45 | 168.36 |
| Building | 346.56 | 609.77 | 248.64 | 231.38 | 180.83 | 305.39 | 77.44 | **352.13\*** | 255.76 | **281.73\*** | 191.17 | **157.75\*** |
| Agriculture | 349.89 | 625.41 | 237.47 | 244.6 | 180.25 | 303.19 | 80.15 | 362.44 | 264.66 | 313.7 | 189.9 | 168.79 |
| Natural | 341.78 | 618.91 | 236.46 | 226.38 | 172.86 | 277.19 | 77.44 | 363.57 | **250.69\*** | 311.44 | **157.83\*** | 161.25 |
| Ag + Nat +<br>Build | **337.58\*** | 607.53 | **231.35\*** | **205.51\*** | **168.46\*** | **273.55\*** | **74.51\*** | **351.51\*** | 255.29 | 291.22 | **157.72\*** | **157.3\*** |
| Rd + Build<br>+ Nat | **338.52\*** | **597.76\*** | 238.29 | **206.6\*** | **167.11\*** | **273.97\*** | **73.41\*** | **351.47\*** | 254.87 | **282.67\*** | **157.91\*** | **156.44\*** |
| Rd + Ag +<br>Nat | 343.87 | 612.42 | **232.19\*** | 224.75 | 170.85 | 275.56 | 76.53 | 360.74 | 263.23 | 288.14 | **157.69\*** | 162.56 |

**Supp. Table 8.** Mean daily displacement (MDD), movement distance (MMD), and movement probabilities (mv prob) calculated for each individual during each season as well as overall

| ID | Cold MDD<br>(m) | Hot MDD<br>(m) | Wet MDD (m) | MDD (m) | Cold mv<br>prob | Hot mv<br>prob | Wet mv<br>prob | Mv<br>prob |
| --- | --- | --- | --- | --- | --- | --- | --- | --- |
| M01 | na | na | 38.35 ± 10.53 | 38.35 ± 10.53 | na | na | 0.34 | 0.34 |
| M02 | 24.13 ± 6.36 | 16.51 ± 4.45 | 67.2 ± 27.64 | 23.17 ± 4.20 | 0.27 | 0.2 | 0.8 | 0.27 |
| M07 | 29.61 ± 8.02 | 4.64 ± 4.42 | na | 24.11 ± 6.38 | 0.22 | 0.04 | na | 0.18 |
| M12 | 33.71 ± 21.45 | 22.73 ± 5.57 | na | 27.01 ± 9.0 | 0.11 | 0.19 | na | 0.16 |
| M14 | 20.42 ± 7.5 | 22.07 ± 9.24 | na | 21.03 ± 5.8 | 0.23 | 0.21 | na | 0.22 |
| F16 | 11.56 ± 9.01 | 4.67 ± 1.77 | na | 5.68 ± 2.0 | 0.13 | 0.12 | na | 0.12 |
| M22 | na | 22.61 ± 5.87 | 60.47 ± 22.72 | 37.0 ± 9.6 | na | 0.48 | 0.58 | 0.52 |
| M27 | na | na | 19.62 ± 10.11 | 19.62 ± 10.11 | na | na | 0.24 | 0.24 |
| M28 | 0 | na | 23.39 ± 5.08 | 19.22 ± 4.26 | 0 | na | 0.35 | 0.29 |
| M29 | na | na | 25.0 ± 22.65 | 25.0 ± 22.65 | na | na | 0.27 | 0.27 |
| M32 | 42.44 ± 15.25 | na | 302.14 ± 111.85 | 60.99 ± 17.37 | 0.19 | na | 0.86 | 0.23 |
| M33 | 15.81 ± 3.65 | 6.27 ± 3.12 | 61.17 ± 39.48 | 14.54 ± 2.98 | 0.19 | 0.1 | 0.5 | 0.18 |
| M35 | 26.94 ± 6.66 | 0 | na | 20.59 ± 5.17 | 0.17 | 0 | na | 0.13 |
| M36 | 60.55 ± 19.38 | 0 | na | 55.98 ± 18.04 | 0.33 | 0 | na | 0.3 |
| *Male Means* | *28.71 ± 3.56* | *14.78 ± 2.1* | *41.34 ± 6.28* | *27.48 ± 2.36* | *0.21* | *0.18* | *0.38* | *0.23* |

